# Impaired Fetal Lung Development can be Rescued by Administration of Extracellular Vesicles Derived from Amniotic Fluid Stem Cells

**DOI:** 10.1101/2020.08.07.240408

**Authors:** Lina Antounians, Vincenzo D. Catania, Louise Montalva, Benjamin D. Liu, Huayun Hou, Cadia Chan, Andreea C. Matei, Areti Tzanetakis, Bo Li, Rebeca Lopes Figueira, Karina Miura da Costa, Amy P. Wong, Robert Mitchell, Anna L. David, Ketan Patel, Paolo De Coppi, Lourenço Sbragia Neto, Michael D. Wilson, Janet Rossant, Augusto Zani

## Abstract

Incomplete lung development, also known as pulmonary hypoplasia, is a recognized cause of neonatal death and poor outcome for survivors. To date, there is no effective treatment that promotes fetal lung growth and maturation. Herein, we describe a novel stem cell-based approach that enhances fetal lung development via the administration of extracellular vesicles (EVs) derived from amniotic fluid stem cells (AFSCs). In experimental models of pulmonary hypoplasia, administration of AFSC-EVs promoted lung branching morphogenesis and alveolarization, and stimulated pulmonary epithelial cell and fibroblast differentiation. This regenerative ability was confirmed in two models of injured human lung cells, where human AFSC-EVs obtained following good manufacturing practices restored pulmonary epithelial homeostasis. AFSC-EV beneficial effects were exerted via the release of RNA cargo, primarily miRNAs, that regulate the expression of genes involved in fetal lung development. Our findings suggest that AFSC-EVs hold regenerative ability for underdeveloped fetal lungs, demonstrating potential for therapeutic application.

**One Sentence Summary:** Fetal lung regeneration via administration of extracellular vesicles derived from amniotic fluid stem cells

## Introduction

Fetal lung development is a crucial step during embryogenesis, which if disrupted leads to a condition called pulmonary hypoplasia. Hypoplastic lungs have a reduced number of bronchiolar divisions, enlargement of airspaces, defective alveolarization, and impaired tissue maturation (*1*). Pulmonary hypoplasia can be idiopathic or secondary to associated malformations, the most common of which is congenital diaphragmatic hernia (CDH) (*1, 2*). CDH is a birth defect characterized by an incomplete closure of the diaphragm that leads to the herniation of intra-abdominal organs into the chest, resulting in pulmonary hypoplasia (*1, 2*). Pulmonary hypoplasia secondary to CDH has a mortality rate of 40% with most babies dying within the first days of life (*3*), and with 60% of survivors suffering from long-term morbidity (*4, 5*). There is an unmet clinical need for an effective treatment that would rescue lung growth and maturation, but to date, none of the therapies tested has been successful (*6*). As pulmonary hypoplasia can be diagnosed as early as at the anatomy scan (18-20 weeks of gestational age), the paradigm of treatment in the last decades has focused on promoting lung growth and maturation before birth (*6*). Lung development is a complex process that is regulated by a network of signaling molecules, including small RNA species. In particular, some miRNAs are known to control biological processes that are important for lung development, such as branching morphogenesis and epithelial and mesenchymal differentiation (*7-9*), and have been reported to be missing or dysregulated in human and animal hypoplastic lungs (*10-13*). Correcting the dysregulated network of signaling molecules would be beneficial to promote lung regeneration in fetuses with pulmonary hypoplasia.

A promising strategy to deliver a heterogeneous population of small RNA species is by administering extracellular vesicles (EVs) (*14-16*). EVs are small, subcellular, biological membrane-bound nanoparticles that carry cargo in the form of genetic material and bioactive proteins (*17-19*). EVs are recognized key mediators of stem cell paracrine signaling and have been shown to promote tissue maturation and regeneration (*20-22*). Amniotic fluid stem cells (AFSCs) could be the ideal source of EVs to promote lung regeneration as AFSCs can integrate and differentiate into epithelial lung lineages (*23*), reduce lung fibrosis (*24*), repair damaged alveolar epithelial cells (*25*), and promote lung growth in a model of pulmonary hypoplasia secondary to CDH (*26, 27*). AFSCs confer regenerative ability despite a low engraftment rate, thus suggesting a paracrine effect (*26-28*), which could, at least in part, be mediated by EVs. Recently, EVs derived from AFSCs have been reported to hold regenerative potential in several animal models, including lung, kidney, and muscle injury *(29)*.

In the present study, we have investigated whether administration of AFSC-EVs to various models of pulmonary hypoplasia could promote growth and maturation of underdeveloped fetal lungs.

## Results

### AFSC-EV administration promotes growth and maturation in fetal hypoplastic lungs

The most robust experimental model of pulmonary hypoplasia relies on nitrofen administration to pregnant rats at embryonic day (E) 9.5 (*30, 31*), which mainly targets retinoic acid synthesis (*32*). In this model, the whole litter has an impairment in lung development that is analogous to that of human fetuses (*2, 30-33*). *In utero* nitrofen exposure causes a reduction in bronchiolar divisions and in the number of airspaces compared to lungs from unexposed fetuses (Fig. 1, A to C) (*30*). Administration of rat AFSC-EVs (mean size 140±5nm) to hypoplastic lung explants harvested during the pseudoglandular stage (E14.5) resulted in an increase in terminal branching morphogenesis (Fig. 1, A to C, fig. S1). Particularly, the total bud count and mean surface area of lung explants treated with AFSC-EVs was increased compared to untreated hypoplastic lungs and similar to control lungs. This effect was specifically due to AFSC-EVs, as administration of AFSC conditioned medium (CM) or EV-depleted AFSC-CM did not rescue the impaired terminal branching. As larger AFSC-EVs (mean size 363±17nm) did not restore branching morphogenesis (fig. S1F), in all the experiments performed hereafter we used small AFSC-EVs (mean size 140±5nm). We also tested whether the effects obtained with AFSC-EVs could be replicated by another source of EVs. There is growing evidence that EVs from mesenchymal stromal cells (MSCs) ameliorate experimental bronchopulmonary dysplasia (*34*), a neonatal lung condition similar to pulmonary hypoplasia secondary to CDH. However, administration of MSC-EVs to hypoplastic lung explants did not restore normal terminal branching (Fig. 1, A to C, fig. S1). We also observed that AFSC-EVs did not affect branching morphogenesis in uninjured control lungs (fig. S2A and B). When we investigated pathways responsible for lung branching morphogenesis, we observed that hypoplastic lungs had lower expression levels of *Fgf10, Vegfα* and its receptors (*Flt1* and *Kdr*), compared to untreated lungs (Fig. 1D) (*35, 36*). AFSC-EV administration improved the expression levels of these factors and receptors (Fig. 1D).

**Fig. 1.**
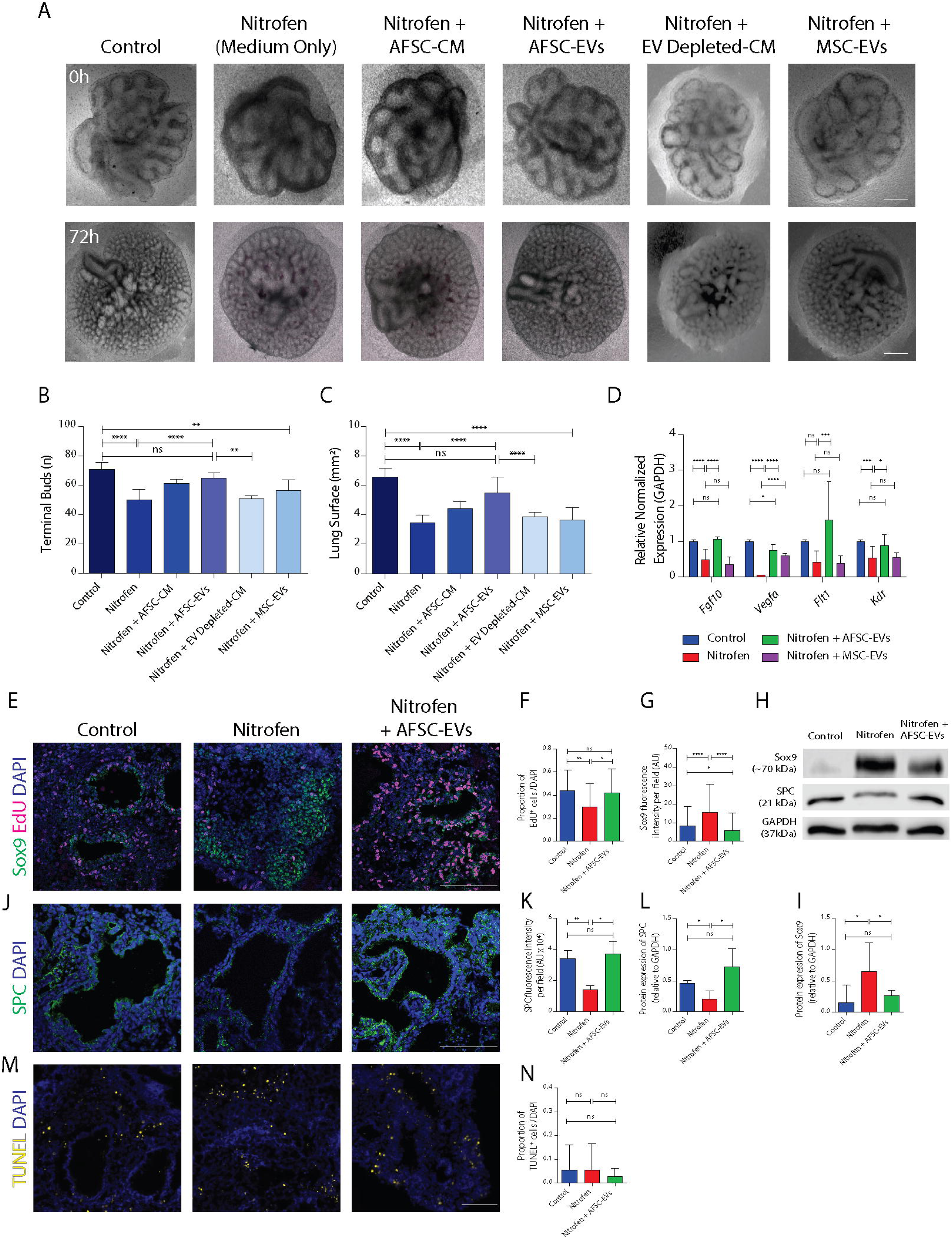
Administration of rat AFSC-EVs promotes growth, branching morphogenesis, and maturation in fetal hypoplastic lungs. (**A**) Representative light microscopy photos of lung explants harvested at E14.5 (0 h) and grown for 72 h in conditions indicated in columns. Scale bar = 750 µm. (**B** and **C**) quantification of total terminal bud count and total lung surface area measured at 72 h. ****P<0.0001, **P<0.01, ns= P>0.05. Data are quantified blindly by two investigators for the following number of biological replicates: Control (n=16), Nitrofen (n=17), Nitrofen+AFSC-CM (n=12), Nitrofen+AFSC-EVs (n=12), Nitrofen+EV-depleted AFSC-CM (n=5), Nitrofen+MSC-EVs (n=8). (**D**) Gene expression changes in lung maturation markers fibroblast growth factor 10 (*Fgf10*) and vascular endothelial growth factor (*Vegfa*), and its receptors (*Flt1* and *Kdr*) in lung explants measured at 72 h. *P<0.05, ***P<0.001. Expression values are averaged between n=3 biological replicates of each condition. (**E** to **G**) Immunofluorescence co-stain experiment of proliferating cells and distal lung epithelium progenitor cells of lung explants (SOX9, green; EdU, pink, DAPI nuclear stain, blue; scale bar = 100 µm), quantified through number of EdU+ cell per DAPI and SOX9 fluorescence intensity (AU= arbitrary units). N=4 biological replicates were used with a total of 50×50 µm fields covering entire lung sections as indicated: Control (n=157), Nitrofen (n=222), Nitrofen+AFSC-EVs (n=128). (**H** and **I**) Western blot analysis of SOX9 and SPC expression in lung explants grown for 72 h, quantified by signal intensity normalized to GAPDH in n=3 biological replicates from each condition. (**J** and **K**) Immunofluorescence experiment of surfactant protein C (SPC) expressing cells in lung explants (SPC, green; DAPI nuclear stain, blue; scale bar =100µm), quantified by fluorescence intensity: Control (n=6), Nitrofen (n=4), Nitrofen+AFSC-EVs (n=4). (**L**) Quantification of SPC protein expression by signal intensity normalized to GAPDH in n=3 biological replicates from each condition. (**M** and **N**) TUNEL immunofluorescence experiments on lung explants grown for 72 h, quantified by TUNEL^+^ cells per DAPI in n=4 biological replicates with a total of 50×50 µm fields covering entire lung sections as indicated: Control (n=311), Nitrofen (n=240), Nitrofen+AFSC-EVs (n=107). Groups were compared using Kruskal-Wallis (post-hoc Dunn’s non parametric comparison) test for Fig. 1 B, C, F, G, I, K, L, N, and with one-way ANOVA (Tukey post-test) for Fig. 1 D, according to Gaussian distribution assessed by D’Agostino Pearson omnibus normality test.

In addition to compromised fetal lung growth, pulmonary hypoplasia is characterized by impaired lung maturation, a feature replicated with *in utero* nitrofen exposure (*2, 30*). Hypoplastic lungs have decreased cell proliferation and delayed epithelial cell differentiation, as demonstrated by an increased number of distal progenitor cells (SOX9+) and by a reduced expression of surfactant protein C (SPC) (Fig. 1E to L, fig. S3A and B) (*32, 37, 38*). AFSC-EV administration rescued cell proliferation back to control levels and improved epithelial cell differentiation, as evidenced by a reduction of SOX9+ progenitor cell density (Fig 1E to F) and by an increased expression of SPC (Fig. 1E-L). Hypoplastic lungs had similar levels of apoptosis throughout the explant compared to control, as previously reported (*33, 39*), and AFSC-EV administration did not alter this phenotype (Fig. 1M-N, fig. S3C).

### AFSC-EV administration restores homeostasis and stimulates differentiation of lung epithelial cells

A hallmark of pulmonary hypoplasia secondary to CDH is the impairment in the homeostasis of the respiratory epithelium (*2, 40*). Administration of AFSC-EVs to primary lung epithelial cells isolated from hypoplastic lungs of nitrofen exposed fetuses increased proliferation and reduced cell death back to control levels (Fig. 2, A and B). We confirmed that these effects were specific to AFSC-EVs, as they were not reproduced by the administration of either AFSC-CM, or EV-depleted AFSC-CM. Interestingly, MSC-EVs improved proliferation of primary lung epithelial cells, but failed to reduce cell death back to control levels (fig. S3D and E). We also observed that administration of AFSC-EVs to uninjured control cells did not change their proliferation or cell death rates (fig. S2C and D).

**Fig. 2.**
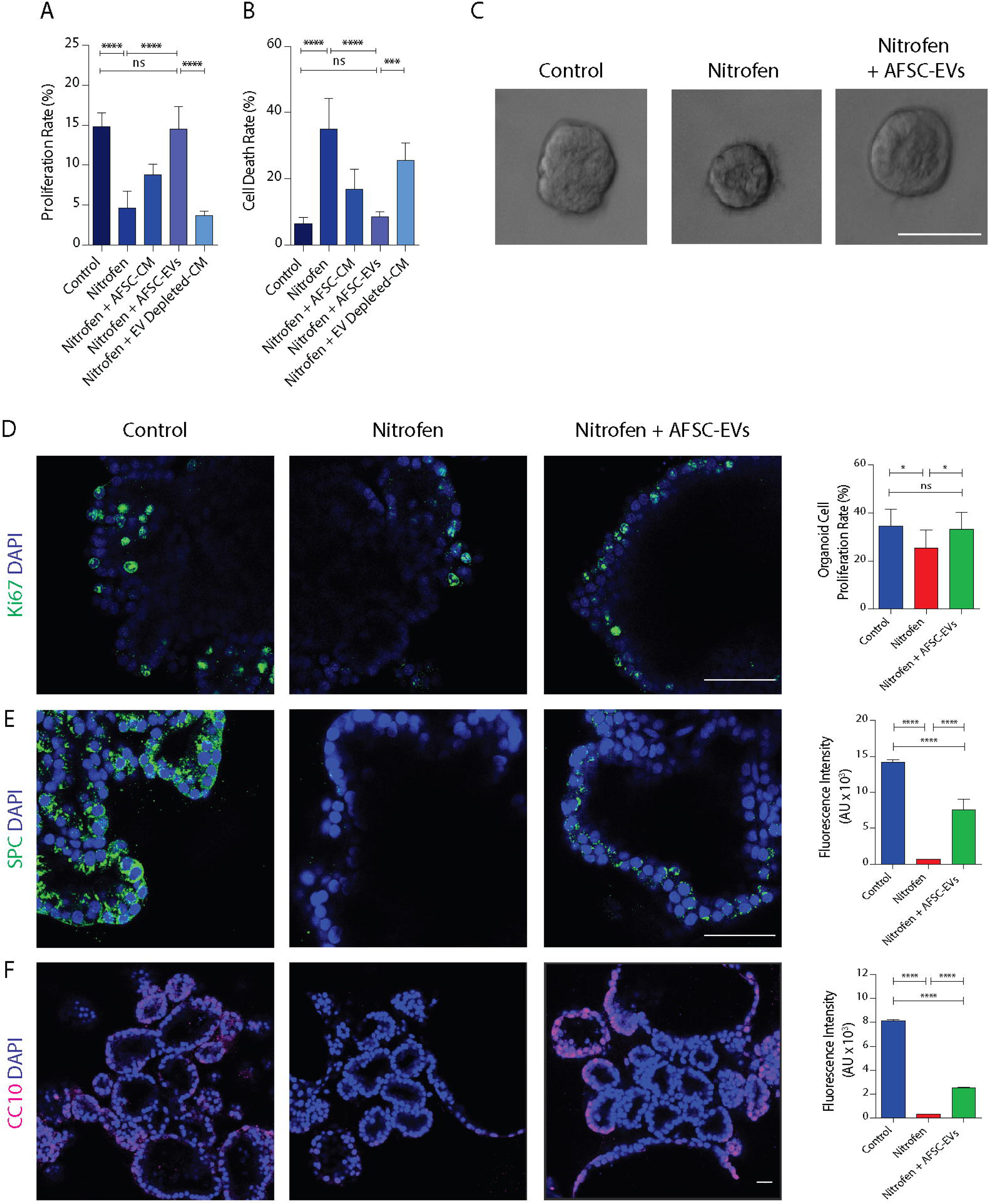
AFSC-EVs maintains homeostasis and stimulate differentiation of the epithelium from hypoplastic fetal lungs. (**A**) Proliferation rate of primary lung epithelial cells from control and nitrofen-exposed hypoplastic lungs treated with AFSC-CM, AFSC-EVs, EV-depleted AFSC-CM [5’EdU labeling, Control (n=7), Nitrofen (n=5), Nitrofen+AFSC-CM (n=3), Nitrofen+AFSC-EVs (n=5), Nitrofen+EV-depleted AFSC-CM (n=3)]. ****P<0.0001, ns = P>0.05. (**B**) Cell death rate of primary lung epithelial cells from control and nitrofen-exposed hypoplastic lungs treated as in (**A**) [live/dead cytotoxicity assay, Control n=5, Nitrofen (n=5), Nitrofen+AFSC-CM (n=6), Nitrofen+AFSC-EVs (n=5), Nitrofen+EV-depleted AFSC-CM (n=4)]. *P<0.05, ***P<0.001. (**C**) Light microscopy photos of fetal rat lung organoids derived from control lungs and nitrofen-exposed hypoplastic lungs either treated with medium alone (Nitrofen) or medium supplemented with AFSC-EVs (Nitrofen + AFSC-EVs). Scale bar = 100 µm. Representative photo of Control (n=108), Nitrofen (n=63), Nitrofen+AFSC-EVs (n=94). (**D**) Proliferation of cells in organoids evaluated with immunofluorescence (Ki67 staining, green; scale bar = 50 µm) and quantified as percentage of Ki67^+^ cells per total number of DAPI (blue) stained nuclei in Control (n=8), Nitrofen (n=7), Nitrofen+AFSC-EVs (n=9). (**E**) SPC staining in organoids (green; DAPI nuclear stain, blue; scale bar = 50 µm) quantified with fluorescence intensity calculated from total corrected cellular fluorescence from Control (n=30), Nitrofen (n=31), Nitrofen+AFSC-EVs (n=25) (AU= arbitrary units). (**F**) CC10^+^ cells in organoids (green; DAPI nuclear stain, blue; scale bar = 50 µm) quantified with fluorescence intensity calculated from total corrected cellular fluorescence Control (n=30), Nitrofen (n=30), Nitrofen+AFSC-EVs (n=30). Groups were compared using Kruskal-Wallis (post-hoc Dunn’s non parametric comparison) test for Fig. 2 A, B, D, E, F according to Gaussian distribution assessed by D’Agostino Pearson omnibus normality test.

To study the effect of AFSC-EVs on respiratory epithelial cell differentiation, we generated fetal lung organoids (Fig. 2C, fig. S3F). The cell proliferation rate of organoids treated with AFSC-EVs was similar to that of organoids derived from control lungs, but higher than that of untreated organoids derived from hypoplastic lungs (Fig. 2D, fig. S3G). Moreover, AFSC-EV treated organoids had a more differentiated respiratory epithelium than untreated organoids, as demonstrated by higher expression levels of SPC (marker of early distal epithelium and alveolar type II cells) and CC10 (marker of club cells) (Fig. 2, E and F, fig. S3H and I).

### AFSC-EV cargo content and its effect on lung epithelium

The beneficial effects of AFSC-EVs on nitrofen exposed hypoplastic lungs and on the respiratory epithelium were associated with the transfer of the EV cargo, which was detected throughout the lung parenchyma (Movie S1) and primary lung epithelial cells (Fig. 3A-B and Movie S2-3). To study the cargo content, we differentially analyzed the proteins and RNA species of AFSC-EVs and MSC-EVs, as we had observed different effects between AFSC-EV and MSC-EV administration to lung explants and primary lung epithelial cells. Proteomics analysis identified 222 differentially expressed proteins, none of which had obvious molecular functions directly related to fetal lung development (fig. S4A). AFSC-EVs contained proteins involved in EV formation (HSPa and CD63) and miRNA stabilization (Annexins and Hnrnps), and proteins involved in pathways important for EV structure and function (table S1).

**Fig. 3.**
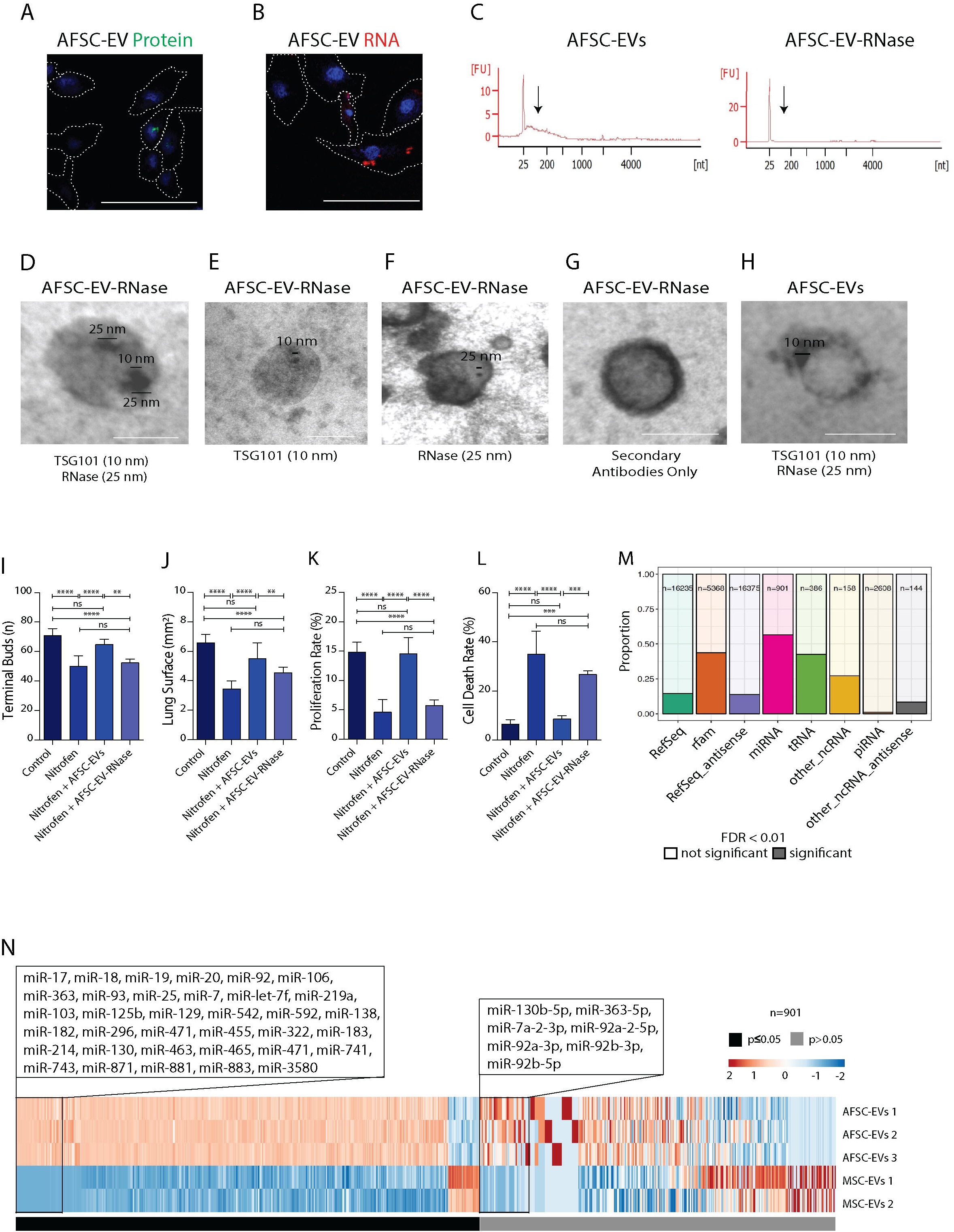
The role of RNA cargo released by AFSC-EVs. (**A** and **B**) Fluorescently labeled AFSC-EV protein (green) and RNA (red) cargo entered primary lung epithelial cells (DAPI nuclear stain, blue; scale bar = 100 µm). Cells were outlined based on light microscopy images to highlight the border. To confirm this, cells were washed twice with PBS and fixed in 4% PFA and re-imaged after live cell imaging (shown in **Movie S2 and Movie S3**). (**C**) Bioanalyzer traces of RNase pre-treated AFSC-EVs shows that RNA cargo content is diminished. (**D** to **H)** Representative photos of gold immunolabeling experiments of RNase-treated AFSC-EVs (**D** to **G**) or AFSC-EVs (**H**), using transmission electron microscopy, with TSG101 (10 nm), and RNase (25 nm). Controls include single stains (**E** and **F**), secondary only antibodies (**G**), and co-stains in untreated AFSC-EVs (**H**). Photos are representative of two biological replicates of AFSC-EV-RNase and AFSC-EVs which included two technical replicates each. (**I** to **L**) Effects of RNase pre-treated AFSC-EVs on lung growth parameters (bud count, (**I**); and surface area, (**J**), from Control (n=16), Nitrofen (n=17), Nitrofen+AFSC-EVs (n=12), and Nitrofen+AFSC-EV-RNase (n=4), and on pulmonary epithelial cells (proliferation, (**K**); and cell death rate (**L**) from Control (n=7), Nitrofen (n=5), Nitrofen+AFSC-EVs (n=5), and Nitrofen+AFSC-EV-RNase (n=4), ****P<0.0001, ***P<0.001, **P<0.01, ns = P>0.05. (**M**) Small RNA-sequencing analysis of AFSC-EVs and MSC-EVs separated by type of RNA species. Solid bar represents proportion of significantly different species per type of RNA. (**N**) Heat map of miRNAs detected in AFSC-EVs (n=3) and MSC-EVs (n=2), ranked based on fold change and significant difference between the two populations. The two inlets report the miRNAs involved in lung development (see table S2). Right inlet: miRNAs significantly enriched in AFSC-EVs detected within the top 50 miRNAs. Left inlet: miRNAs equally abundant in AFSC-EVs and MSC-EVs detected within the top 50 miRNAs. Groups were compared using Kruskal-Wallis (post-hoc Dunn’s non parametric comparison) test for Fig. 3 I, J, K, L according to Gaussian distribution assessed by D’Agostino Pearson omnibus normality test.

To test the role of the RNA cargo on fetal lung development, we performed an RNase-A enzymatic digestion of AFSC-EVs. We verified that RNase-A degraded the RNA cargo content and was captured by AFSC-EVs, as shown with immune-electron microscopy by its co-localization with TSG101 inside the AFSC-EVs (Fig. 3C-H). When we added RNase-pretreated AFSC-EVs to hypoplastic lung explants or to primary epithelial cells derived from hypoplastic lungs, we did not observe an increase in lung terminal branching and surface area, or an increase in cell proliferation and decrease in cell death, as seen with AFSC-EVs (Fig. 3, I-L). We confirmed that this effect was not due to a carry-over effect secondary to the transfer of RNase-A to the epithelial cells (fig. S4B). This suggested that the delivery of AFSC-EV RNA cargo was a mediator of their biological effects on fetal lung development. We next used RNA-sequencing to identify and compare the small RNA cargo between AFSC-EVs and MSC-EVs. We found both AFSC-EV and MSC-EV cargos contained mRNA, tRNA, miRNA, and piRNA (Fig. 3M, fig. S4C). Among all, miRNAs were the RNA species most proportionally different between the cargos of the two populations. Compared to MSC-EVs, AFSC-EVs were enriched for miRNAs that are critical for lung development, such as the miR17∼92 cluster and its paralogues (miRs-93, -106, -250, and -363; fold enrichment ranging from 5.1 - 9.36; table S2, Fig. 3N). Moreover, AFSC-EVs contained miRNAs that have previously been reported as dysregulated in hypoplastic lungs, such as miR-33 and miR-200 (table S3) (*10, 12*).

To identify the regulatory pathways affected by AFSC-EVs, we used mRNA-sequencing to compare gene expression between primary lung epithelial cells from nitrofen-exposed lungs treated with AFSC-EVs or MSC-EVs. We also applied the same treatment on primary lung epithelial cells from normal lungs. Nitrofen-exposure significantly altered the gene expression profile of primary lung epithelial cells compared to uninjured normal control epithelial cells (fig. S4D). Using gene set enrichment analysis, we found that AFSC-EV administration to nitrofen exposed primary epithelial cells altered the expression of genes related to epithelial differentiation and homeostasis maintenance, whereas MSC-EV administration altered the expression of genes involved with cell cycle regulation and nuclear organization (Fig. 4A and B, fig. S5A and B). To understand the effects of AFSC-EVs on fetal lung epithelial cells, we manually queried these genes and identified those that are critical for lung development (table S4). We then asked if the miRNAs identified in the AFSC-EV cargo were part of predicted regulatory networks with the mRNAs that were down-regulated in the target epithelial cells (Fig. 4C). The network that resulted from this analysis showed that there were genes important for lung development and that were regulated by the miR17∼92 cluster and its paralogues. Small RNA-sequencing of the nitrofen exposed primary epithelial lung cells treated with AFSC-EVs showed that most of the miRNAs in the network had higher expression levels compared to the nitrofen only group (Fig. 4C). Lastly, we correlated the miRNA cargo content with the miRNA target cell content, and their validated mRNAs in the target cells (fig. S6). This triple analysis identified likely miRNA-mRNAs pairs that might be responsible for the phenotype observed. This analysis provides indirect evidence that the miRNA cargo content of AFSC-EVs was transferred to the target cells after EV conditioning.

**Fig. 4.**
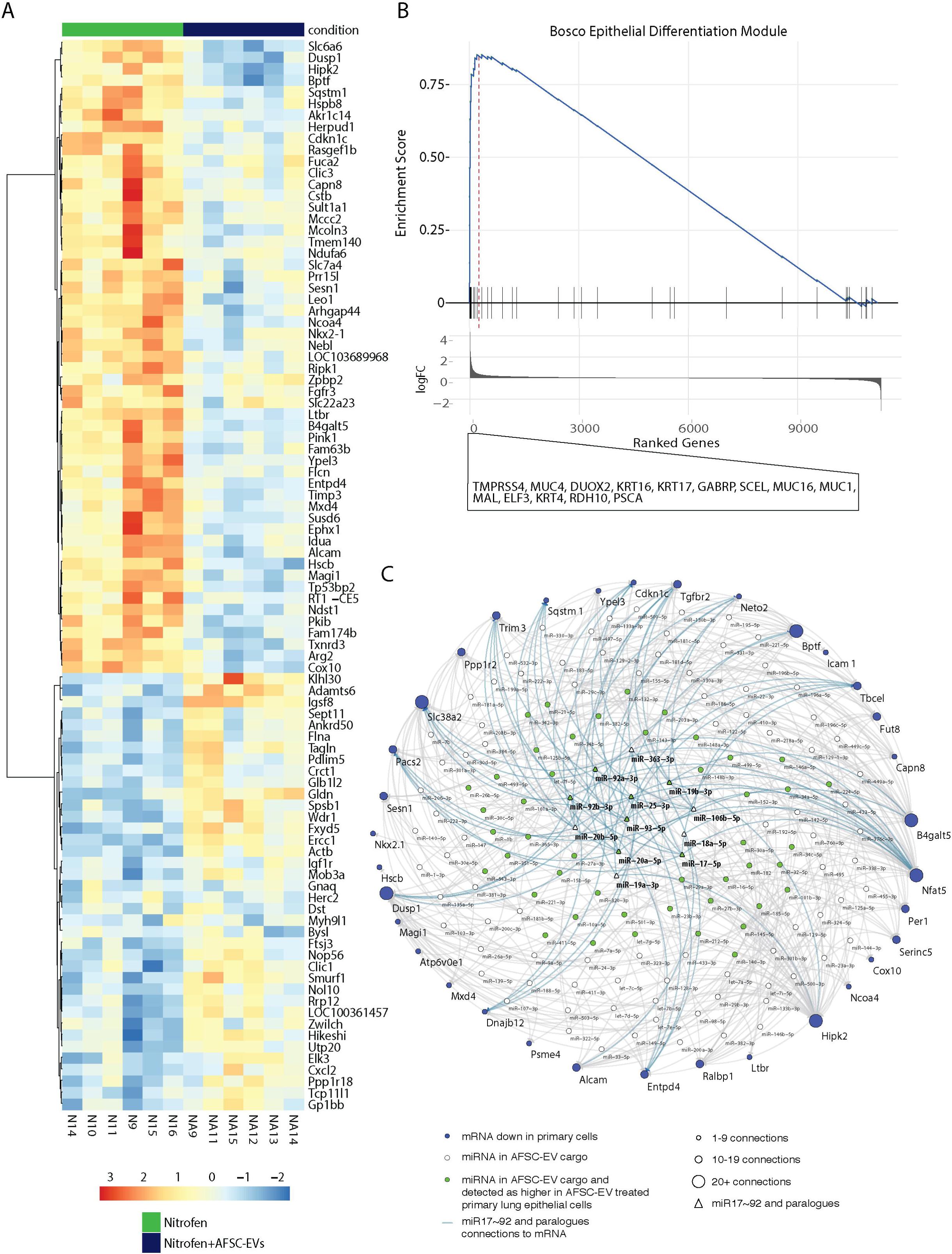
Effects of AFSC-EVs exerted on primary lung epithelial cells. (**A**) Heat map of mRNA expression of genes from lung epithelial cells of nitrofen-exposed lungs treated with AFSC-EVs from n=6 biological replicates each. Only genes with an FDR < 0.1 in the differential analysis are shown. Color represents row-scaled, normalized read counts (RPKM). (**B**) GSEA enrichment plot of epithelial cell differentiation generated with genes ranked by fold change between NA vs. N. “Leading edge” genes are shown on the side. (**C**) Interaction network of AFSC-EV miRNAs and down regulated genes in nitrofen-exposed lung epithelial cells treated with AFSC-EVs. Size of node represents number of connections. Blue nodes represent genes down-regulated (FDR < 0.1) in nitrofen-exposed lung epithelial cells treated with AFSC-EVs compared to nitrofen-exposed only lung epithelial cells. White nodes represent miRNAs that were detected in the AFSC-EV cargo. Green nodes represent miRNAs that were detected in AFSC-EV cargo that had higher expression in nitrofen-exposed lung epithelial cells treated with AFSC-EVs compared to nitrofen-only treated epithelial cells. Triangles represent miRNAs from the miR17∼92 family and paralogues. Each miRNA-gene target pair is connected by a gray edge and pairs containing a miRNA from miR17∼92 family or its paralogues are connected by a blue edge.

### Towards the clinical translation of AFSC-EVs as a treatment for fetal lung regeneration

To test the effects of AFSC-EVs *in vivo*, we used a surgical model of pulmonary hypoplasia and CDH in fetal rabbits, as it allowed us to topically deliver our treatment (Fig. 5A) (*41*). To prevent AFSC-EV egression, we performed tracheal occlusion on rabbit fetuses (Movie S4), a procedure that is currently used in clinical trials for human fetuses with CDH that have the worst cases of pulmonary hypoplasia (*42*). Administration of rat AFSC-EVs improved lung alveolarization by increasing the number of alveoli, decreasing the alveolar wall thickness, and promoting alveolar lipofibroblast maturation (Fig. 5B to E). Moreover, only lungs of fetuses that received AFSC-EVs had improved levels of BMP signaling (BMP2, BMP4, Id1), which is important for alveolar maturation (Fig. 5F). These effects were not observed upon MSC-EV administration (fig. S5C to E).

**Fig. 5.**
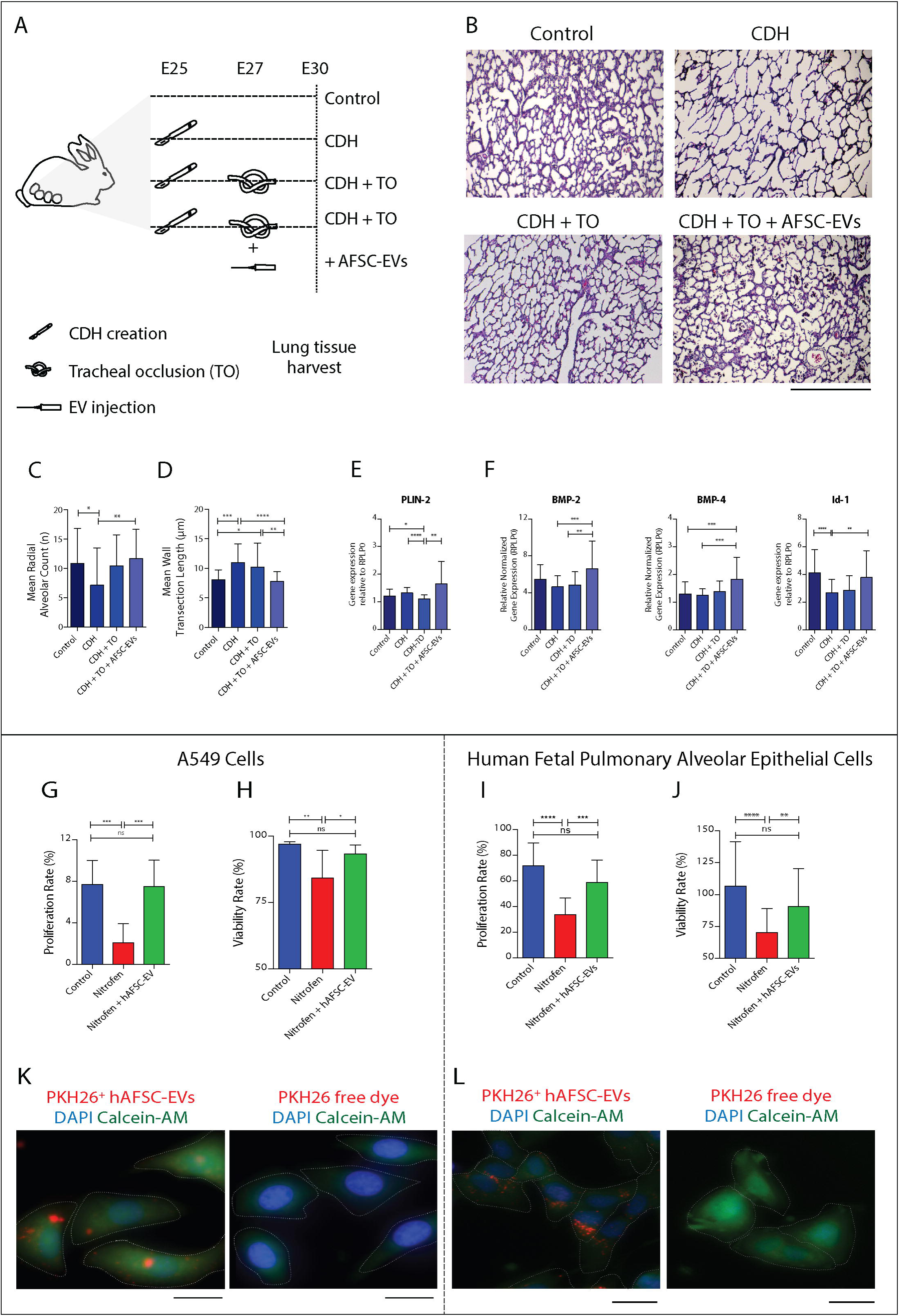
Towards the clinical translation of AFSC-EVs as treatment for fetal lung regeneration. (**A**) Schematic of experimental groups from the rabbit model of CDH. (**B**) Representative histology images (hematoxylin/eosin) of fetal lungs from control rabbits and from rabbits that underwent surgical CDH creation and were either untreated (CDH), or subjected to tracheal occlusion (CDH+TO), or were administered rat AFSC-EVs prior to tracheal occlusion (CDH+TO+AFSC-EVs). Each condition included fetal lungs from n=9 experiments. Scale bar = 500 µm. (**C** and **D**) Differences in number of alveoli (radial alveolar count) were quantified in at least 12 counts per fetal lung in 2 different sections of the lung, and thickness of the alveolar wall (mean wall transection length) was measured in 10 different areas of each lung. **P<0.01, *P<0.05. (**E** and **F**) Gene expression changes in alveolar lipofibroblasts (PLIN2), BMP (bone morphogenetic protein) signaling (BMP2, BMP4, Id1) in n=9 biological replicates of each condition. ***P<0.001 (**G** and **J**) Effects of good manufacturing practice-grade human AFSC-EVs on proliferation rate and viability rate on nitrofen injured human A549 cells (**G** and **H**), and human pulmonary alveolar epithelial cells (**I** and **J**) conducted in n=3 technical replicates. (**K** and **L**) Uptake of human AFSC-EVs fluorescently labeled with PKH26 by A549 cells or human pulmonary alveolar epithelial cells. Scale bar = 25 µm. Groups were compared using Kruskal-Wallis (post-hoc Dunn’s non parametric comparison) test for Fig. 5 C, D, E, F, H, I, and J, and with one-way ANOVA (Tukey post-test) for Fig. 5 G according to Gaussian distribution assessed by D’Agostino Pearson omnibus normality test.

To investigate whether AFSC-EV beneficial effects could be translated onto human lung tissue, we obtained AFSCs from donated human amniotic fluid following clinically compliant good manufacturing practices (GMP), as previously reported (*22*). Administration of human AFSC-EVs replicated similar effects on cellular homeostasis observed in the rat model on both a validated model of lung injury with nitrofen exposed human alveolar epithelial (A549) cells and on nitrofen exposed primary human pulmonary alveolar epithelial cells (HPAEpiC) isolated from a fetus at 21 weeks of gestation (Fig. 5G to J). We verified that fluorescently labeled PKH26+ human AFSC-EVs entered A549 cells and HPAEpiC (Fig. 5K and L). Administration of human MSC-EVs to HPAEpiC did not rescue cell homeostasis, despite entering the cells (fig. S7A to C). Lastly, when we tested human AFSC-EVs on the *in vivo* rabbit model, we found similar effects on alveolar wall thickness, as found with rat AFSC-EV administration (fig. S7D to F).

## Discussion

In this study, we have shown for the first time that administration of AFSC-EVs to various models of pulmonary hypoplasia promotes fetal lung regeneration. Specifically, AFSC-EVs administered to fetal hypoplastic lungs rescued branching morphogenesis and alveolarization by promoting epithelial and mesenchymal tissue growth and maturation and by re-establishing cellular homeostasis (Fig. 6). These beneficial effects were obtained through the release of AFSC-EV cargo. In particular, we identified miRNAs contained in AFSC-EVs that regulate lung maturation processes.

**Fig. 6.**
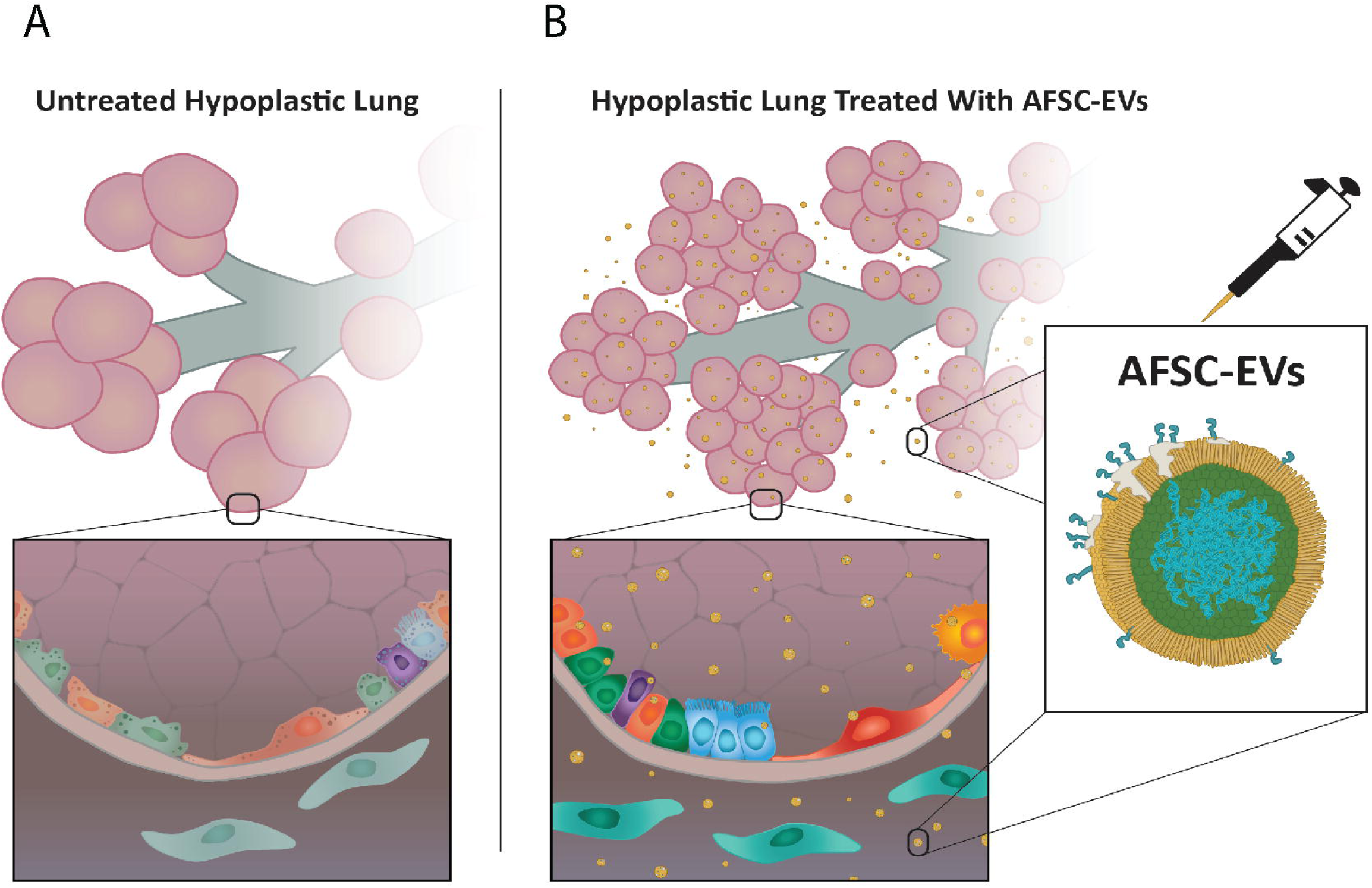
Schematic of changes that occur with AFSC-EV administration to hypoplastic fetal lungs. (**A**) Untreated hypoplastic lungs with few lung buds, small total surface area, and dysregulated homeostasis (low cell proliferation, high cell death, undifferentiated epithelial and fibroblast cells). (**B**) Administration of AFSC-EVs to hypoplastic lungs improves terminal branching and results in increased number of lung buds, higher surface area, and restored cellular homeostasis (improved cell proliferation, reduced cell death, more differentiated epithelial and fibroblast cells).

EVs are emerging as a successful strategy to promote tissue maturation and regeneration in various models. Their regenerative potential that we have observed in our models of fetal pulmonary hypoplasia has been observed in other models of tissue regeneration using either AFSC-EVs (*29*) or EVs from other stem cell sources (*14-16, 20, 43, 44*). In fact, there is increasing evidence that stem cell secreted EVs carry cargo that stimulates typical stem cells-like paracrine functions on target cells such as renewal, differentiation, and maturation (*45-47*). In this study, we confirmed that AFSC-EVs have a similar effect on hypoplastic lung explant and on lung epithelium to that exerted by their parent cells, AFSCs (fig. S8A to D). For this reason and because they are considered immunologically innocuous, EVs provide an advantageous and safer cell-free alternative compared to a stem cell-based therapy (*48-50*). Although conditioned medium could also be obtained in a GMP fashion and be a potential therapy in humans (*51-53*), in our study the EV fraction of the CM was more potent than the whole CM or than the EV-depleted CM fraction, as also observed by others (*54, 55*). Possibly, this effect is due to the fact that EVs carry active molecules that are more concentrated than in their parent cells (fig. S8E). This is in line with our previous findings reporting that AFSC-EV concentration is the most important parameter responsible for their regenerative potential (*29*). Likewise, in the present study, we have shown that the impact of AFSC-EVs is dependent on the concentration of EVs isolated from the CM, as well as on the size of the vesicles that enter the target cells (fig. S1D-G, Movie S2-3, and S5). How size contributes to biological function of EVs remains unknown (*56*). Nonetheless, small EVs, also called exosomes (*17*), have traditionally been considered the EV subpopulation with more potent, protective, and pathological functions than larger vesicles, and therefore with more potential as diagnostic or therapeutic tool (*29*).

We used MSCs as an alternative source of stem cell EVs, as these cells are currently being tested in several clinical trials for the treatment of bronchopulmonary dysplasia (*57*), the postnatal lung condition that is most comparable to pulmonary hypoplasia. In our study, MSC-EV administration to our pulmonary hypoplasia models did not have similar beneficial effects as AFSC-EV administration, despite entering the primary lung cells (fig. S2-3, Movie S6). The different effects obtained with the two populations of EVs may be due to differences in the disease pathogenesis, where pulmonary hypoplasia is mainly the result of abnormal and delayed lung development, and bronchopulmonary dysplasia is a chronic postnatal lung disease with substantial pulmonary inflammatory response (*58*). Moreover, our analysis of the EV cargo identified profile differences between the two EV populations, which could explain the outcome differences with MSC-EV administration.

In our models of pulmonary hypoplasia, we identified that the RNA cargo of AFSC-EVs was key to regenerate hypoplastic fetal lungs. This finding is in line with the knowledge that EV-mediated effects occur through the transfer of RNA species (*14-17*). We found that miRNAs were the RNA species most differentially represented in the cargo between AFSC-EVs and MSC-EVs. When considering the AFSC-EV specific genes in the context of miRNA cargo enriched in AFSC-EVs, we found miRNA species that were important for the phenotypes that we observed, including branching morphogenesis, alveolarization, homeostasis, and cell differentiation (table S2). Specifically, a family of miRNAs that is enriched in AFSC-EVs compared to MSC-EVs is the miRNA 17∼92 cluster. This cluster is key for lung branching morphogenesis (*59*), and when knocked out causes severe fetal pulmonary hypoplasia, making this a candidate mechanism that warrants further investigation (*9*). Moreover, the small RNA sequencing analysis revealed that members of this cluster were up-regulated in the nitrofen exposed primary lung epithelial cells following treatment with AFSC-EVs (fig. S8F).

An improvement in fetal lung maturation was observed not just when the same species of AFSC-EVs were administered on the same species of target cells (i.e. rat AFSC-EVs on rat lung tissue or human AFSC-EVs on human lung epithelial cells), but also when we tested rat AFSC-EVs or human AFSC-EVs on the *in vivo* rabbit model (Fig. 4A to F, fig. S7D to F). We speculate that the improvement in alveolarization observed in rabbit fetuses can be explained by the fact that some miRNAs and their targets are conserved across species (*60*). For instance, from the top 50 miRNAs that were enriched in the human AFSC-EVs, 13 are evolutionarily conserved between human and rabbit species, including one member of the miR17∼92 cluster member (table S5).

Our study provides insights into the potential use of AFSC-EVs as a therapy for fetal pulmonary hypoplasia. Using GMP-grade human AFSC-EVs, we have confirmed similar beneficial effects on damaged epithelial cells derived from a fetal lung at the gestational age when pulmonary hypoplasia and CDH are typically detected. Further steps are needed for AFSC-EVs to be used as a therapy in humans. One of the challenges will be identifying the most effective and safest route of administration to fetuses. In this study, we have employed topical administration via intra-tracheal injection in rabbit fetuses. This route could be further explored, also in clinical settings, as it is currently used to occlude the trachea of human fetuses with CDH that have the worst cases of pulmonary hypoplasia (*42*).

We acknowledge that our study has some limitations. Our findings are mainly based on the use of animal models and human lung epithelial cells to study a complex human condition with an unknown etiology. However, obtaining human lung tissues from babies with pulmonary hypoplasia is not considered ethically acceptable and nor has it been reported. Moreover, in this study we have focused mainly on the analysis of the fetal lung epithelium and mesenchyme, but we have not examined other lung tissue types potentially affected by AFSC-EV administration, such as pulmonary vessels, which are known to undergo vascular remodeling in hypoplastic lungs. Nonetheless, we have observed improvements in pathways important for lung vascular development, such as VEGF and FGF10 that suggest the opportunity for further studies. Another limitation is that little is known about factors that alter lung development and the mechanisms that are dysregulated in pulmonary hypoplasia. Similarly, it remains also unknown how exactly the EV RNA cargo species function. It is hoped that our transcriptomic profiling of both hypoplastic lungs and EVs will increase the understanding on the pathogenesis of pulmonary hypoplasia and on EV function.

## Materials and Methods

### Study Design

The objective of this study was to evaluate the ability of AFSC-EV administration to promote fetal lung growth and maturation in pulmonary hypoplasia. As obtaining human lung tissues from babies with CDH is not considered ethically acceptable, part of this study was conducted using animal models that closely resemble the degree of pulmonary hypoplasia that is observed in human fetuses. To advance towards a translational therapy, we obtained EVs from GMP-grade human AFSCs isolated from donated amniotic fluid during amniocentesis. We tested these human AFSC-EVs first onto a validated model of lung injury with the use of A549 alveolar epithelial cells (*29, 61*). To more closely replicate the conditions of a human fetal lung, we also investigated the effects of GMP-grade human AFSC-EVs on nitrofen-exposed human pulmonary alveolar epithelial cells obtained from a healthy fetus at 21-weeks of gestation. As detailed below, experimental models and sample collections were approved by the appropriate regulatory committees at: The Hospital for Sick Children, Toronto, ON, Canada (AUP#39168 and 49892); University College London Hospital, London, UK (UCL/UCLH Joint Committee for the Ethics of Human Research, REC Reference: 08/0304); Ribeirão Preto Medical School, University of São Paulo, Ribeirão Preto, São Paolo, Brazil (191/2018, 40/2020). Sprague-Dawley rats and New Zealand rabbits were used for all animal studies. Samples from all models were randomly assigned to treatment groups. All data including outliers is shown, and all experiments were performed in at least triplicate, with the number of replicates indicated in the figure legends. Analysis of data was conducted by two blinded investigators. Additional details on the methods used in this study are provided in the Supplementary Materials.

### EV isolation, characterization, and tracking

EVs from rat and human AFSCs or bone marrow MSCs were isolated by ultracentrifugation from conditioned medium of cells that were treated with exosome-depleted FBS for 18 h, as previously described (*29*). In accordance with the International Society for Extracellular Vesicles guidelines, AFSC-EVs and MSC-EVs were characterized for size using Nanoparticle Tracking Analysis, morphology by transmission electron microscopy, and expression of canonical EV-related protein markers by Western blot analysis, as previously described (*29*). To track EV migration into primary lung epithelial cells and lung explants, AFSC-EV and/or MSC-EV cargo was fluorescently labelled for RNA and protein using Exo-Glow™, and for lipid membrane using PKH26 red fluorescent cell linker. RNase enzymatic digestion of AFSC-EVs was conducted in a subset of experiments using AFSC-EVs to determine the role of RNA in fetal lung explants and cell cultures. A bioanalyzer was used to confirm the efficacy of RNase digestion. Co-labelled TSG101 and RNase were visualized by immuno-electron microscopy.

### Experimental models of pulmonary hypoplasia

*Ex vivo -* In fetal rats, pulmonary hypoplasia was induced as previously described (*30, 40*) with the administration of nitrofen to pregnant dams (100mg) by oral gavage on E 9.5. At E14.5, the dam was euthanized, and fetal lungs were micro-dissected. Lungs were grown as explants on nanofilter membranes for 72 h in culture medium alone (DMEM), AFSC-CM, AFSC-EVs (10% by volume), or MSC-EVs (10% by volume). Fetal lungs from dams that received olive oil (no nitrofen) at E9.5 served as control.

*In vitro* – 1) For primary epithelial cell experiments in rats, a single cell suspension was obtained at E14.5 from pooled lungs from either control or nitrofen exposed rat fetuses by trypsinization for 20 minutes. Cells were subjected to three serial depletions of fibroblasts by incubation for 1h each following an established protocol (*40, 62*). 2) For organoid studies, cells were seeded in a ratio of 60:40 semi-solid Matrigel to medium, as previously described (*63*). Cells from nitrofen-treated fetuses were cultured for 10 days with medium alone or with medium supplemented with 10% v/v AFSC-EVs or MSC-EVs. Lung organoids from fetuses whose mothers had not received nitrofen served as control. 3) A549 cells were treated for 24h with nitrofen (40µM), and treated with medium alone, human AFSC-EVs (10% v/v), or human MSC-EVs (10% v/v). Uninjured and untreated A549 cells served as control. 4) Human pulmonary alveolar epithelial cells (HPAEpiC) were obtained from the lungs of a healthy fetus at 21 weeks of gestation and used at first passage. Cells were stressed with 400µM nitrofen for 24h, and then treated with medium alone, or medium supplemented with 10% v/v human AFSC-EVs or human MSC-EVs.

*In vivo -* In fetal rabbits, pulmonary hypoplasia was induced secondary to surgical creation of a diaphragmatic hernia at E25 (*41*) in New Zealand rabbits. At E27, tracheal ligation was performed either alone or after intra-tracheally administration of a 50µL bolus containing either rat AFSC-EVs, rat MSC-EVs, or human AFSC-EVs (Movie S4). Lungs were harvested at E31 and immediately frozen for RNA extractions or fixed in 4% paraformaldehyde and embedded in paraffin.

### Outcome measures

For lung morphometry, rat fetal lung explants were imaged by differential interference contrast microscopy and independently assessed by two blinded researchers for terminal bud density and surface area using ImageJ, as previously described (*27*). Rabbit fetal lungs were blindly evaluated with histology (hematoxylin and eosin staining) to assess the number of alveoli and the thickness of the alveolar wall, as described (*64, 65*). For RNA expression, factors involved in rat or rabbit lung branching morphogenesis were assessed with quantitative polymerase chain reaction (RT-qPCR). For studies on lung tissue homeostasis on lung explants and organoids, 5-ethynyl-2’-deoxyuridine (EdU incorporation kit (Click-IT®), LIVE/DEAD™ Viability/Cytotoxicity kit, Terminal deoxynucleotidyl transferase dUTP nick end labeling (TUNEL assay - Click-IT®), and Ki67+ immunofluorescence staining were used. For studies on epithelial differentiation, lung explants were immunostained for SOX9 and SPC, assessed for expression of SPC, a marker of differentiated epithelial cells. To determine the main effectors of AFSC-EVs, the RNA cargo content was sequenced (small RNA-seq) and organoids for SPC and CC10. To study the AFSC-EV protein cargo, we used nanoscale liquid chromatography coupled to tandem mass spectrometry (nano LC-MS/MS), and conducted differential expression analysis using 1% protein and peptide false discovery rate (FDR). For the RNA cargo, we conducted small RNA-sequencing on rat AFSC-EVs and MSC-EVs and compared RNA content using Bioconductor DESeq. To study the effects of AFSC-EV RNA cargo on target cells, total RNA was isolated from primary lung epithelial cells (n=6 replicates) from each condition (Control, Nitrofen, Nitrofen+AFSC-EVs, and Nitrofen+MSC-EVs) and edgeR was conducted for comparative analysis. To study changes in miRNA expression patterns in target cells, we further analyzed the Nitrofen and Nitrofen+AFSC-EVs treated cells (n=3-4 replicates) described above using small RNA sequencing. edgeR was used to determine miRNA expression changes (FDR<0.1), and Spearman’s correlation was used to assess potential gene targets of detected miRNAs in cells. Pathway enrichment analysis was conducted with g:profiler and miRNA-mRNA network analysis was performed with TargetScan, miRTarBase and Cytoscape.

### Statistical Analysis

Groups were compared using two-tailed Student t-test, Mann-Whitney test, one-way ANOVA (Tukey post-test), or Kruskal-Wallis (post-hoc Dunn’s non-parametric comparison) test according to Gaussian distribution assessed by D’Agostino Pearson omnibus normality test. For correlation studies, a Pearson coefficient was reported as r (confidence interval). P value was considered significant when p<0.05. All statistical analyses were produced using GraphPad Prism^®^ software version 6.0.

mRNA and miRNA sequencing analyses in lung epithelial cells were performed in R (version 3.6.0). Package “edgeR” (version 3.26.5) was used for differential analyses between two conditions with FDR < 0.1 considered as significant for both mRNA and small RNA-seq.

## Supplementary Materials

Fig. S1. Characterization of AFSC-EVs and MSC-EVs and effects on lung explants based on size and concentration.

Fig. S2. Effects of AFSC-EVs on control lung explants and primary lung epithelial cells.

Fig. S3. Effects of MSC-EV administration on *in vitro* and *ex vivo* models of pulmonary hypoplasia.

Fig. S4. Analysis of protein and RNA cargo of AFSC-EV and MSC-EV.

Fig. S5. Enrichment plots for RNA-seq analysis of MSC-EV treated primary lung epithelial cells and effects on *in vivo* model of pulmonary hypoplasia.

Fig. S6. Correlation analysis of miRNA-mRNA sequencing in primary lung epithelial cells.

Fig. S7. Effects of human AFSC-EVs and human MSC-EVs on human fetal lung epithelial cells and in the *in vivo* model of pulmonary hypoplasia.

Fig. S8. Influence of AFSCs in co-culture with *ex vivo* and *in vitro* models of pulmonary hypoplasia and effects on AFSC-EV miRNA cargo.

Table S1. Highlighted proteins expressed in AFSC-EVs and MSC-EVs.

Table S2. miRNAs related to lung development that are differentially expressed in AFSC-EVs over MSC-EVs.

Table S3. miRNAs known to be involved in pulmonary hypoplasia and present in AFSC-EVs.

Table S4. Genes differentially expressed in nitrofen-exposed lung epithelial cells.

Table S5. Inter-species conservation of top enriched miRNA in human AFSC-EVs

Table S6: Primer sequences used in this study.

Table S7. Details of antibodies used in this study.

Movie S1. AFSC-EV tracking into lung explant.

Movie S2. Live tracking of AFSC small EV RNA into lung epithelial cells.

Movie S3. Live tracking of AFSC small EV Protein into lung epithelial cells.

Movie S4. *In vivo* administration of AFSC-EVs prior to tracheal ligation in fetal rabbits.

Movie S5. Live tracking of AFSC medium/large EV RNA into lung epithelial cells.

Movie S6. Live tracking of MSC small EV RNA into lung epithelial cells.

Data file S1 (Microsoft Excel format). Differential analysis of AFSC-EV and MSC-EV protein cargo using proteomic analysis.

Data file S2 (Microsoft Excel format). Quality control table for all RNA-sequencing experiments used in this study.

Data file S3 (Microsoft Excel format). Full list of differentially expressed genes with pair-wise comparisons between Nitrofen vs. Nitrofen+AFSC-EVs, and Nitrofen vs. Nitrofen+MSC-EVs.

Data file S4 (Microsoft Excel format). Full list of pathway enrichments for mRNA-sequencing experiments.

## Acknowledgments

The authors would like to thank Reta Aram, Alyssa Belfiore, George Biouss, Dr. Jennifer Guadagno, Sasha Korogodski, Dr. Kasra Khalaj, Dr. Yuhki Koike, Dr. Kimberly Lau, Carol Lee, Dorothy Lee, Qi Ma, Karim Maghraby, Dr. Ornella Pellerito, Dr. Gabriele Raffler, Mark Stasiewicz, Dr. Adrienne Sulistyo, Dr. Yanting Wang, and Dr. Kyoko Yuki. Moreover, the authors would thank Dr. Christian Smith and Dr. James Rutka from the Brain Tumour Research Centre, the Lab Animal Service core facility and Nanoscale Biomedical Imaging Facility at the Hospital for Sick Children.

## Author contributions

L.A., V.D.C., L.S.N., M.D.W., J.R., and AZ designed the research study. L.A., V.D.C., L.M., B.D.L., A.C.M., A.T., B.L., R.L.F., K.M.C., and L.S.N. performed the in vitro and in vivo experiments, collected and analyzed the data. A.P.W. supervised the organoid studies. R.M., A.L.D., K.P., and P.D.C. isolated and characterized human amniotic fluid stem cells. L.A., H.H., C.C., M.D.W., and AZ analyzed bioinformatics data. L.A., A.Z. wrote the manuscript. P.D.C., A.P.W., M.D.W., J.R. provided critical reading of the manuscript. All authors approved the final manuscript.

## Data and materials availability

Proteomics differential analysis with AFSC-EV and MSC-EV cargos can be found in Supporting Data File S1. We have submitted all relevant data of our EV experiments to the EV-TRACK knowledgebase (EV-TRACK ID: EV190001). RNA-sequencing data for AFSC-EVs, MSC-EVs, and primary lung epithelial cells from all experimental groups are available in the ArrayExpress database (http://www.ebi.ac.uk/arrayexpress).

## Figure Legends

**Fig. S1. Characterization of AFSC-EVs and MSC-EVs and effects on lung explants based on size and concentration.** (**A**) Representative plot of the average size distribution of AFSC-EVs and MSC-EVs visualized using Nanoparticle Tracking Analysis. Data are representative of five 40-second videos of each EV preparation. X-axis = size distribution (nm), y-axis = concentration (particles/mL). (**B**) Representative transmission electron microscopy photos of AFSC-EVs and MSCEVs; two different magnifications highlight the morphology of individual EVs at near fields (top) and far fields (bottom). Scale bar = 200 nm. (**C**) Expression of canonical EV markers TSG101, Flo-1, Hsp70, and CD63 obtained by Western blot analysis for AFSC-EVs and MSC-EVs in n=3 technical replicates. EV preparations do not express histone marker H3K27me3, indicating preparations are free of cell debris. (**D**) Representative plot of the average size distribution of medium/large AFSC-EVs. Data are representative of five 40-second videos of each EV preparation. X-axis = size distribution (nm), y-axis = concentration (particles/mL). (**E**) Live cell tracking of RNA in medium/large AFSC-EVs (m/lAFSC-EVs), DAPI, blue. Scale bar = 100 µm. Cells were outlined based on light microscopy images to highlight the border. (**F**) Effects of EV size (small EVs, AFSC-sEVs; medium/large, m/lAFSC-EVs) on lung explant terminal bud count and mean surface area in Control (n=16), Nitrofen (n=17), Nitrofen+AFSC-CM (n=12), Nitrofen+AFSC-sEVs (n=12), and AFSC-m/lEVs (n=4). (**G**) Effects of decreasing doses of AFSC-EVs (10%, 5%, and 1%) on lung explant terminal bud count and mean surface area in Control (n=16), Nitrofen (n=17), Nitrofen+10% AFSC-EVs (n=12), Nitrofen+5% AFSC-EVs (n=5), Nitrofen+1% AFSC-EVs (n=5). Groups were compared using Kruskal-Wallis (post-hoc Dunn’s non parametric comparison) test for fig. S1 F and G, according to Gaussian distribution assessed by D’Agostino Pearson omnibus normality test.

**Fig. S2. Effects of AFSC-EVs on control lung explants and primary lung epithelial cells.** (**A** and **B**) Addition of AFSC-EVs on control lung explants quantified for terminal bud count, (**A**) and lung surface area, (**B**), in Control (n=16) and Control+AFSC-EVs (n=4). Effects on control primary lung epithelial cells proliferation (**C**), and cell death rates (**D**) in Control (n=4) and Control+AFSC-EVs (n=3). ns = P>0.05. Groups were compared using Mann-Whitney test for fig. S2 A-D, according to Gaussian distribution assessed by D’Agostino Pearson omnibus normality test.

**Fig. S3. Effects of MSC-EV administration on *in vitro* and *ex vivo* models of pulmonary hypoplasia.** (**A**) Immunofluorescence co-stain experiment of proliferating cells and distal lung epithelium progenitor cells of lung explants (SOX9, green; EdU, pink, DAPI nuclear stain, blue; scale bar = 100 µm), quantified through number of EdU+ cell per DAPI and SOX9 fluorescence intensity (AU= arbitrary units) in n=4 biological replicates with a total of 50×50µm fields covering entire lung sections as indicated: Control (n=157), Nitrofen (n=222), Nitrofen+AFSC-EVs (n=128), Nitrofen+MSC-EVs (n=122). (**B**) Immunofluorescence experiment of surfactant protein C (SPC) expressing cells in lung explants (SPC, green; DAPI nuclear stain, blue; scale bar=100µm), quantified by fluorescence intensity: Control (n=6), Nitrofen (n=4), Nitrofen+AFSC-EVs (n=4), Nitrofen+MSC-EVs (n=4). (**C**) TUNEL immunofluorescence experiments on lung explants grown for 72 h, quantified by TUNEL+ cells per DAPI in n=4 biological replicates with a total of 50×50µm fields covering entire lung sections as indicated: Control (n=311), Nitrofen (n=240), Nitrofen+AFSC-EVs (n=107), Nitrofen+MSC-EVs (n=191). (**D**) Proliferation rate of primary lung epithelial cells from control and nitrofen-exposed hypoplastic lungs treated with medium only, AFSC-EVs, or MSC-EVs [5’EdU labeling Control (n=7), Nitrofen (n=5), Nitrofen+AFSC-EVs (n=5), Nitrofen+MSC-EVs (n=4)]. (**E**) Cell death rate of primary lung epithelial cells from control and nitrofen-exposed hypoplastic lungs treated as in (**D**) (live/dead cytotoxicity assay in Control n=5, Nitrofen (n=5), Nitrofen+AFSC-EVs (n=5), Nitrofen+MSC-EVs (n=4). **(F)** Light microscopy photos of fetal rat lung organoids derived from nitrofen-exposed hypoplastic lungs treated with MSC-EVs. Scale bar = 100 µm. Representative photo of n=104 organoids imaged. (**G**) Proliferation of cells in organoids evaluated with immunofluorescence (Ki67 staining, green; scale bar = 50 µm) and quantified as percentage of Ki67^+^ cells per total number of DAPI (blue) stained nuclei in Control (n=8), Nitrofen (n=7), Nitrofen+AFSC-EVs (n=9), Nitrofen+MSC-EVs (n=14). (**H**) SPC staining in organoids (green; DAPI nuclear stain, blue; scale bar = 50 µm) quantified with fluorescence intensity calculated from total corrected cellular fluorescence from Control (n=30), Nitrofen (n=31), Nitrofen+AFSC-EVs (n=25), Nitrofen+MSC-EVs (n=5). (**I**) CC10^+^ cells in organoids (green; DAPI nuclear stain, blue; scale bar = 50 µm) quantified with fluorescence intensity calculated from total corrected cellular fluorescence from Control (n=30), Nitrofen (n=30), Nitrofen+AFSC-EVs (n=30), Nitrofen+MSC-EVs (n=25). Groups were compared using Kruskal-Wallis (post-hoc Dunn’s non parametric comparison) test for fig. S3 A-E, G-I, according to Gaussian distribution assessed by D’Agostino Pearson omnibus normality test.

**Fig. S4. Analysis of protein and RNA cargo of AFSC-EV and MSC-EV. (A)** Pathway enrichment analysis of AFSC-EV enriched protein cargo in GO terms Biology Processes, Cellular Component, and Molecular Functions. (**B**) Effect of potential carry-over of RNase in AFSC-EV-RNase experiments on control cell proliferation in n=3 biological replicates. The enzymatically treated AFSC-EVs were separated from the supernatant, which was then administered to control lung epithelial cells. Groups were compared using unpaired t-test, according to Gaussian distribution assessed by D’Agostino Pearson omnibus normality test. (**C**) Heatmap showing expression levels of significantly different species of RNA (FDR<0.01) separated by type for cargo from AFSC-EV (n=3) and MSC-EV (n=2) samples. Rows and columns are displayed using hierarchical clustering (Ward’s method; row distance measure: Pearson correlation; column distance measure: Euclidean). Color scale represents rows scaled (scaled across samples for each gene), log2 transformed normalized counts. (**D**) Volcano plot of genes differentially expressed between nitrofen-exposed lung epithelial cells and control (non-nitrofen exposed) lung epithelial cells. Each dot represents a gene with log2 fold change <-2 (lower expression in nitrofen vs. control, n=322 red nodes) or >2 (higher expression in nitrofen vs. control, n=661 blue nodes).

**Fig. S5. Enrichment plots for RNA-seq analysis of MSC-EV treated primary lung epithelial cells and effects on *in vivo* model of pulmonary hypoplasia.** (**A**) Scatterplot of log fold changes between Nitrofen+AFSC-EVs (NA) vs. Nitrofen (N) and Nitrofen+MSC-EVs (NM) vs. Nitrofen. Pearson correlation = 0.54 (p < 2.2e-16, 95% CI [0.53, 0.55]). Each dot represents a gene with its log fold change between NA vs. N shown on the X-axis and log fold change between NM vs. N shown on the Y-axis. Dots are colored based on if the gene is identified as significantly differentially expressed (FDR < 0.1) only in NA vs. N (red), in NM vs. N (blue), or in both comparisons (purple). See supplementary data file S3 for the full lists of differentially expressed genes. (**B**) Selected pathways identified with gene set enrichment analysis of NA vs. N and NM vs. N primary lung epithelial cells for GO pathways Biological Process, Cellular Component, and Molecular Function. See supplementary data file S4 for full lists of GSEA results. (**C**) GSEA enrichment plot of epithelial cell differentiation generated with all genes ranked by fold changes between Nitrofen exposed epithelial cells vs. Nitrofen + MSC-EV-treated. (**D**) Representative histology images (hematoxylin/eosin) of fetal lungs from fetal rabbits that underwent surgical CDH creation and were subjected to administration of MSC-EVs prior to tracheal occlusion (CDH+TO+MSC-EVs). Scale bar = 500 µm. (**E** and **F**) Differences in number of alveoli (radial alveolar count, **E**) and thickness of the alveolar wall (mean wall transection length, **F**) between CDH+TO+AFSC-EVs (n=9) and CDH+TO+MSC-EVs (n=8). Groups were compared using Mann-Whitney test for fig. S5 E and F, according to Gaussian distribution assessed by D’Agostino Pearson omnibus normality test.

**Fig. S6. Correlation analysis of miRNA-mRNA sequencing in primary lung epithelial cells.** Negatively correlated miRNA-mRNA pairs in primary lung epithelial cells are graphed by log of counts per million mapped reads (CPM) of miRNA (purple) and mRNA (orange) for primary lung epithelial cells from nitrofen (N) and nitrofen+AFSC-EVs (NA) conditions. For a given negatively correlated pair (n=3-4 replicates), logCPM expression is shown for miRNA (purple) and gene (orange). Expression is shown for nitrofen-exposed only lung epithelial cells (N, circles) and nitrofen-exposed+AFSC-EVs lung epithelial cells (NA, triangle). Spearman’s correlation coefficient (rho) is shown next to miRNA-mRNA pair label.

**Fig. S7. Effects of human AFSC-EVs and human MSC-EVs on human fetal lung epithelial cells and in the *in vivo* model of pulmonary hypoplasia.** (**A**) Proliferation rate of primary human pulmonary alveolar epithelial cells (HPAEpiC) from a fetus at 21 weeks of gestation, injured with 400µM of nitrofen exposure for 24h, and then treated with medium only, or medium supplemented with GMP-grade human AFSC-EVs or human MSC-EVs. Replicates were conducted in n=4 technical replicates. (**B**) Number of viable cells per field for the same conditions as in (**A**). (**C**) Human MSC-EVs labeled with PKH26 (red signal) entered HPAEpiC (live cells labeled with Calcein-AM, green, nucleus labeled with DAPI, blue). Scale bar = 25 µm. (**D**) Representative histology image (hematoxylin/eosin) of fetal lungs rabbits that underwent surgical CDH creation and were administered human AFSC-EVs prior to tracheal occlusion (CDH + TO + human AFSC-EVs). Scale bar = 500 µm. Samples are representative of Control (n=9), CDH (n=9), CDH+TO (n=9), CDH+TO+AFSC-EVs (n=9), and CDH+TO+hAFSC-EVs (n=5). (**E** and **F**) Differences in number of alveoli (radial alveolar count, **E**) and thickness of the alveolar wall (mean wall transection length, **F**). Groups were compared using Kruskal-Wallis (post-hoc Dunn’s non parametric comparison) test for fig. S7 E, F, according to Gaussian distribution assessed by D’Agostino Pearson omnibus normality test.

**Fig. S8. Influence of AFSCs in co-culture with *ex vivo* and *in vitro* models of pulmonary hypoplasia and effects on AFSC-EV miRNA cargo.** (**A** and **B**) Effects on terminal bud count (A) and mean surface area (B) of nitrofen-exposed fetal lung explants co-cultured with AFSCs. Explants were grown on top of AFSCs for 72 h. The data are from n=7 different fetal lung explants co-cultured with AFSCs. (**C** and **D**) Proliferation rate (**C**) and cell death rate (**D**) of primary lung epithelial cells treated in co-culture with AFSCs in transwells for 24h. The data are from n=3 biological replicates of co-cultured cells. (**E**) miRNA expression of AFSCs compared with AFSC-EVs through miRCURY LNA SYBR qPCR. Data are shown as normalized relative expression to MSCs and MSC-EVs for four members of the miR17∼92 family (n=3 matched replicates per each condition). (**F**) Log of counts per million mapped reads (CPM) for the expression of miR17∼92 family members and paralogues in primary lung epithelial cells exposed to nitrofen (N, n=4, grey triangles) and nitrofen+AFSC-EVs (NA, n=3, pink circles) identified through small RNA-sequencing analysis. Groups were compared using Kruskal-Wallis (post-hoc Dunn’s non parametric comparison) test for fig. S8 A and B, and with Mann-Whitney test for fig. S8 C and D, according to Gaussian distribution assessed by D’Agostino Pearson omnibus normality test. Groups in fig. S8 E were compared using Holm-Sidak method with alpha=5%, each target was analyzed individually, without assuming a consistent SD.

