## Supplementary Methods and Tables for "Impaired Fetal Lung Development can be Rescued by Administration of Extracellular Vesicles Derived from Amniotic Fluid Stem Cells"

**One Sentence Summary:** Fetal lung regeneration via administration of extracellular vesicles derived from amniotic fluid stem cells

***Extended Materials and Methods***

*Cells*

Rat amniotic fluid stem cells (AFSCs) were isolated from amniotic fluid of E12 Sprague-Dawley rat fetuses as previously described (*66*), and grown in alpha-minimal Essential Media (αMEM, Gibco, ThermoFisher, Waltham, MA) supplemented with 20% Chang supplements (Irvine Scientific, Santa Ana, CA), 15% fetal bovine serum (FBS, ThermoFisher Scientific, Waltham, MA), and 0.5% Penicillin/Streptomycin (ThermoFisher Scientific, Waltham, MA). Human AFSCs were obtained under good manufacturing practice guidelines as previously described (UCL/UCLH REC Reference: 08/0304) (*22*). Sprague-Dawley rat bone-marrow derived mesenchymal stem cells (MSCs) were purchased (CellBiologics, Chicago, IL) and grown in supplier recommended medium until 90% confluence. Human adenocarcinomic alveolar basal epithelial cells (A549) were purchased (Sigma Aldrich, Missouri, MO) and grown in DMEM F-12 media (Gibco, ThermoFisher, Waltham, MA) supplemented with 10% FBS and 0.5% Penicillin/Streptomycin. All cells were used within 6 passages. Human pulmonary alveolar epithelial cells (HPAEpiC) were obtained from the lungs of a healthy fetus at 21 weeks of gestation (ScienCell, Carlsbad, CA), grown in supplier recommended medium, and used immediately for proliferation and viability assays. Human bone marrow-derived MSCs were obtained from a healthy donor (ATCC, Manassas, Virginia) and grown in supplier recommended media (Mesenchymal Stem Cell Growth Kit, PCS-500-041). Human MSCs were used within three passages.

*Extracellular vesicles (EVs)*

EVs from rat and human AFSCs and from MSCs were isolated from cells that were treated with exosome-depleted FBS (ThermoFisher, Waltham, MA) for 18 h by ultracentrifugation as previously described (*29*). EVs used in this study were characterized for nanoparticle size and morphology, and expression of canonical EV-related markers (fig. S4), as previously described (*29*). Small EVs had a mean size of 140±5 nm and mode size 104±11 nm). Nanoparticle tracking analysis was conducted on n=3 samples with n=8 40-second videos captured by the NanoSight LM10 system. Data was collected, averaged, and analyzed using LM10 NTA equipped with a 65 mWatt 405 nm violet laser. For size calibration, 100 nm polystyrene beads were used. At capture with sCMOS camera on NTA 3.1, Build 3.1.46, the temperature was 22 °C.

To isolate medium/large EVs (m/lEVs, >200nm), sucrose gradient ultracentrifugation was used with six layers ranging from 10% to 90% sucrose in PBS. AFSC-CM was spun through this gradient at 100,000g for 14h. Fractionated layers were isolated and nanoparticle tracking analysis was used to confirm the presence of large vesicles (m/lEVs mean size of 363±17 nm, mode size 217±32 nm).

*Ex vivo model of pulmonary hypoplasia*

In fetal rats, pulmonary hypoplasia was induced (#39168, #49892) as previously described (*30, 40*) with the administration of nitrofen (Sigma Aldrich, Missouri, MO) to pregnant Sprague-Dawley rats (100 mg in 1 ml olive oil). At E14.5, the dam was euthanized, and fetal lungs were harvested. Lungs were washed in phosphate buffered saline (PBS; Wisent, Saint-Jean-Baptiste, QC), grown on nanofilter membranes (Whatmann Nucleopore membranes, Sigma Aldrich, Missouri, MO), and incubated for 72 h in culture medium alone (DMEM), AFSC-CM, AFSC EVs (400 μg/ml), or MSC EVs (400 μg/ml). For AFSC co-culture experiments, five thousand AFSCs were seeded onto the bottom of the plate, and lung explants on nanofilter membranes were added on top of the cells and allowed to float for 72 h. Fetal lungs from dams that received olive oil (no nitrofen) at E9.5 served as control.

*In vitro model of pulmonary hypoplasia*

At E14.5, a single cell suspension was obtained from pooled lungs from either control or nitrofen exposed rat fetuses by trypsinization (0.25% Trypsin-EDTA, ThermoFisher, Waltham, MA) for 20 minutes. Cells were spun down by centrifugation (5 minutes, 800g) and the pellet was resuspended in DMEM supplemented with 10% FBS and subjected to three serial depletion of fibroblasts by incubation for 1 h each following an established protocol (*40, 62*). Cells were used to test epithelial homeostasis and to generate fetal lung organoids for testing epithelial differentiation. For epithelial homeostasis experiments, cells were grown for 5 days in Bronchial Epithelial Cell Growth Medium (BEGM; Lonza, Basel Switzerland). Cells were checked daily for proper epithelial morphology using a light microscope. On the fifth day, cells were confirmed to be positive for SPC and negative for vimentin via immunofluorescence staining assays. Epithelial homeostasis was investigated by assessing cell proliferation and cell death as previously described (*40*) on cells from nitrofen-exposed lungs had either their medium replaced with BEGM, or BEGM supplemented with 500uL of AFSC-CM, 200μg/mL of AFSC-EVs, EV-depleted AFSC-CM, or 200μg/mL of MSC-EVs. For AFSC co-culture experiments, AFSCs were seeded in the top compartment of a transwell (0.4μm) and primary lung epithelial cells were seeded in the bottom compartment (in a ratio of 1:10, AFSC to primary lung epithelial cells), as previously described (*40*).

For epithelial differentiation experiments, cells were seeded in a ratio of 60:40 semi-solid Matrigel (Corning, Corning, NY) to media ratio, to generate organoids as previously described (*63*). Cells from nitrofen-treated fetuses were cultured for 10 days with medium alone or with medium supplemented with 10% by volume AFSC-EVs or MSC-EVs. Lung organoids from untreated fetuses served as control. Medium was replaced every other day.

Human A549 cells were treated for 24h with nitrofen (40μM), then subsequently administered medium alone, human AFSC-EVs (10% by volume), or human MSC-EVs (10% by volume). Untreated and uninjured A549 cells served as control. Following 24h incubation, proliferation and cell death rates were determined by EdU incorporation kit (Click-IT®, ThermoFisher Scientific, Waltham, MA), and Live/Dead cell viability assay (ThermoFisher Scientific, Waltham, MA). Experiments were repeated as many times as indicated: control (n=4), medium only (n=4), human AFSC-EVs (n=4), human MSC-EVs (n=4) per each assay.

HPAEpiC from a fetus at 21 weeks of gestation contained both alveolar type I and type II cells, which were selected with cytokeratin-18 and cytokeratin-19 by the supplier. Cells were grown in supplier recommended medium and used in the first passage after reaching ~70% confluence. HPAEpiC were stressed with nitrofen exposure at 400μm (dosage at 40μm was not optimal in inducing impairment in cell proliferation or viability). Experiments were repeated as many times as indicated: control (n=6), medium only (n=6), human AFSC-EVs (n=4), human MSC-EVs (n=4) per each assay.

*In vivo model of pulmonary hypoplasia*

In fetal rabbits, pulmonary hypoplasia was induced secondary to surgical creation of a diaphragmatic hernia at embryonic day E25 in New Zealand rabbits, following ethical approval (191-2018), as previously described *(41)*. Two days later at E27, tracheal ligation was performed either alone or in conjunction with EV administration [rat AFSC-EVs (n=9), rat MSC-EVs (n=8), or human AFSC-EVs (n=5)]. EVs (50uL) were injected intra-tracheally prior to ligation of the trachea as shown in Movie S4. Lungs were harvested at E31 and immediately frozen for RNA extractions or were fixed in 4% paraformaldehyde and embedded in paraffin. Fetal rabbits with intact diaphragms and that did not receive tracheal occlusion served as control.

*Lung morphometry*

In fetal rats, lung explants from different conditions were compared for terminal bud density and surface area using ImageJ independently by two blinded researchers, as previously described (*27*). Terminal branching was measured by counting the number of terminal buds defined as the number of single acini separated by distinct septae at the periphery of the explant (*27*). Differential interference contrast photos were taken on a light microscope (Leica DMI6000B, Wetzlar, Germany) at 2.5X magnification. Experiments were repeated as many times as indicated: control (n=16), medium only (n=17), AFSC-CM (n=12), 10% AFSC-EVs (n=12), EV-depleted AFSC-CM (n=5), RNase-treated AFSC-EVs (n=4), MSC-EVs (n=8), 5% AFSC-EVs (n=5), 1% AFSC-EVs (n=5), AFSC- m/lEVs (n=4), and co-cultured AFSCs (n=7).

Rabbit fetal lungs were blindly evaluated with histology (H&E staining) to assess the number of alveoli and the thickness of the alveolar wall, as measures of the degree of lung alveolarization. For the number of alveoli, the radial alveolar count (RAC), a well-established index of alveolar number within the acinus (*64*), was determined in at least 12 counts per fetal lung in 2 different sections of the lung. For the thickness of the alveolar wall, the mean wall transection length was measured in 10 different areas of each lung, as previously described (*65*). Each condition included fetal lungs from n=9 experiments, except MSC-EV-treated group contained n=8 pups, and human AFSC-EV-treated group contained n=5 pups.

*RNA expression*

Lung explants from fetal rats and freshly harvested lungs of fetal rabbits were frozen at -20°C. Total RNA was isolated using Trizol reagent following supplier recommended protocols (ThermoFisher Scientific, Waltham, MA). Purified RNA was quantified using a NanoDrop™ spectrophotometer (ThermoFisher Scientific, Waltham, MA) and cDNA synthesis was performed with 200ng for rat and 1µg for rabbit quantified RNA (superscript VILO cDNA synthesis kit, ThermoFisher Scientific, Waltham, MA). qPCR experiments were conducted with SYBR™ Green Master Mix (Wisent, Saint-Jean-Baptiste, QC) for 40 cycles (denaturation: 95°C, annealing: 58°C, extension: 72°C) using the primer sequences reported in table S6. Melt curve plots were used to determine target specificity of the primers. ΔΔCT method was used to determine normalized relative gene expression.

To determine the expression of specific miRNA in EVs and parent cells, RNA was extracted from approximately 4 million cells of rat AFSCs and rat MSCs (n=3 replicates each) using the Nucleospin miRNA kit (Macherey-Nagel, Düren, Germany), following supplier recommended protocols. EV RNA was isolated using the same procedure, from CM of the corresponding parent cells (n=3 replicates each). cDNA was synthesized using miRCURY LNA RT kit (Qiagen, Hilden, Germany) and UniSp6 RNA Spike-in controls were used. Expression of miR-17-5p (YP02119304), miR-18a-5p (YP00204207), miR-19b-3p (YP00204450), miR-20a-5p (YP00204292), and controls Rnu5g (YP00203908) was assessed with miRCURY LNA SYBR® Green PCR Kit (Qiagen, Hilden, Germany), following supplier recommended protocols. 40-cycle qPCR was conducted as described above, and ΔΔCT method was used to determine normalized relative miRNA expression of AFSC-EVs and AFSCs to MSCs and MSC-EVs.

*Immunofluorescence*

Rat lung explants were fixed using 4% paraformaldehyde (Sigma Aldrich, Missouri, MO) for 20 minutes, washed in PBS, and incubated in 30% sucrose (Sigma Aldrich, Missouri, MO) for at least 18 h. Optimal cutting temperature (Electron Microscopy Sciences, Hatfield, PA) embedded lungs were cryosectioned in coronal orientation and stained with primary antibodies reported in table S7. Fetal lung organoids were fixed in 4% paraformaldehyde for 30 minutes, permeabilized in 0.5% triton X (Sigma Aldrich, Missouri, MO) and 0.05% Tween-20 (Sigma Aldrich, Missouri, MO) in PBS, pre-blocked in 1% bovine serum albumin (BSA; Sigma Aldrich, Missouri, MO) with 0.2% triton-X and 0.05% Tween-20 in PBS. Organoids were stained with primary antibodies reported in table S7. A Leica SP8 lightning confocal microscope (Wetzlar, Germany) was used to image samples using the same laser power and exposure across conditions. Wherever possible, z-stacks were taken to increase coverage of tissues. Total corrected cellular fluorescence was calculated to compare the fluorescence intensities between conditions, as previously described (*67*) using ImageJ 1.51.

*Proliferation and apoptosis experiments on lung explants*

Proliferation experiments in explants were conducted with the addition of EdU (10µM final concentration in media) to lung explants cultures 3 h prior to the 72 h endpoint of experiments. Explants were then fixed and processed for immunofluorescence assays as stated above. The Click-iT™ protocol was followed to label EdU+ cells (2.4:1000, Alexa Fluor™ 647) in co-staining experiments with Sox9 (1:1000, Alexa Fluor™ 488), as recommended by the supplier. Lung explants from n=4 biological replicates were used in at least triplicate technical replicates for analysis. Quantification of EdU signal was conducted with HistoQuant in QuantCenter Imaging Software (3D Histech, Budapest, Hungary).

Cell apoptosis experiments on lung explants were conducted with the Click-iT™ TUNEL assay for in situ apoptosis detection, according to manufacturer's recommended protocol. Briefly, cryo-sectioned lung explants from n=4 biological replicates were used in at least triplicate technical replicates for analysis. Alexa Fluor™ 647 was used in co-staining experiments. Quantification of TUNEL signal was conducted with HistoQuant in QuantCenter Imaging Software (3D Histech, Budapest, Hungary).

*Protein expression*

Protein from lung explants was isolated by re-suspending explants in cell extraction buffer (ThermoFisher Scientific, Waltham, MA) supplemented with protease inhibitors (Sigma Aldrich, Missouri, MO), and sonicating for 3 cycles of 10 seconds each. Protein was quantified using the Pierce Bradford Assay (ThermoFisher Scientific, Waltham, MA), and 20µg of protein from each sample was processed as described (*29*) and probed for SPC and Sox9 (table S7). Expression of canonical EV markers CD63, Hsp70, Flo-1, and TSG101 in AFSC-EVs and MSC-EVs was analyzed as previously described (*29*). All details for the antibodies are available in table S7.

*EV characterization and staining*

To track AFSC-EV migration into primary lung epithelial cells and lung explants, their cargo was fluorescently labelled for RNA and protein using with Exo-Glow™ (System Biosciences, Palo Alto, CA), and for lipid membrane using PKH26 red fluorescent cell linker (Sigma Aldrich, Missouri, MO) following supplier recommended protocols. For PKH26 staining, EVs isolated using ultracentrifugation were re-suspended in Diluent C, stained with PKH26 for five minutes with periodic mixing, then the reaction was stopped with 1% BSA in water. Starting samples containing water only were used as negative controls for staining procedures. For live cell tracking, cells were grown on 35mm µ-Dish plates for 24h to reach 60% confluence (Ibitreat, Fitchburg, WI). DAPI was added in culture media for 10 minutes (2% in media), and then 2µg of EVs stained with ExoGlow RNA or ExoGlow Protein were added and imaged once every two seconds for 10 minutes using a Leica SP8 lightning confocal microscope. After live cell imaging was performed, cells were washed twice with PBS and fixed in 4% PFA, and re-imaged to visualize the internalization of the stained EVs.

*Role of AFSC-EV RNA Cargo*

In selected experiments, EVs were treated with RNase-A (ThermoFisher, Waltham, MA) at 2μg/μL for 90 minutes, then with RNase inhibitor (ThermoFisher, Waltham, MA) as described (*68*). Data shown in viability and proliferation assays are representative of n=3 technical replicates, with at least 5 fields per experiment. Bioanalyzer analysis (Agilent Technologies, Santa Clara, CA) was used to test effectiveness of RNA degradation, after total RNA isolation using miRvana miRNA isolation kit as described above. Untreated EVs that were exposed to all temperature changes without the addition of RNase were used as control.

To confirm the entry of RNase into AFSC-EVs, TSG101 and RNase were probed by immuno-EM labelling. Samples were prepared for TEM following previously established protocols (*29*). Briefly, fixed EV preparations (AFSC-EVs or RNase-treated AFSC-EVs) were allowed to absorb onto charged EM grids for 1 h. Samples were washed in PBS, permeabilized in 0.05% triton-X in PBS for 30 minutes, washed five times in PBS, and then incubated for 2 h in primary antibody (1:100 in 10% BSA-c; Aurion, Wageningen, Netherlands) with TSG101, RNase, or both (table S7). Following fives washes in PBS, samples were incubated for 1 h with the corresponding secondary antibodies (1:25 in 10% BSA-c, table S7), washed an additional five times in PBS, then fixed in 1% glutaraldehyde for 10 minutes. Following ten washes in distilled water, grids were contrasted and embedded in uranyl oxalate for 10 minutes, methyl cellulose-uranyl acetate for 10 minutes, and then prepared for EM imaging on a Tecnai 20 (FEI, Hillsboro, OR) from 25 kx to 100 kx magnification. Experimental groups included single stains, and only secondary antibody stains (incubation with serum alone). Experiments were optimized for incubation time and antibody concentrations. Gold tags were identified and measured manually using the Tecnai 20 Software. To determine if there was a carry-over effect of the RNase treatment on AFSC-EVs, we performed an additional step and separated the enzymatically treated AFSC-EVs from the supernatant, which presumably contained the inactivated RNase. This supernatant was then administered to the control primary lung epithelial cells, and proliferation rate was determined as described above.

*Profiling of AFSC-EVs and MSC-EVs*

10 mL of AFSC- or MSC-CM was centrifuged at 1500g for 5 min to remove residual cells and debris. The supernatant was transferred to a new 50 ml conical tube for EV isolation. Isolations were conducted in triplicate. ExoQuick-TC (System Biosciences, Palo Alto, CA) was added to the supernatant at 1:5 ratio (ExoQuick:Supernatant), mixed gently, and allowed to incubate for 18 h at 4°C. After 24 h, the admixture was centrifuged at 1500 g for 30 min to recover EVs.

*Proteomic profiling of EV cargo*

EVs were quantified for protein concentration using Qubit fluorometry (Invitrogen, Carlsbad, CA). 10µg of EV protein was processed by SDS-PAGE using 10% Bis Tris NuPage mini-gel (Invitrogen, Carlsbad, CA) in the MES buffer system. The migration window (2cm lane) was excised and in-gel digestion was performed using a ProGest robot (DigiLab, Hopkinton, MA) with the following protocol: 1) Washed with 25mM ammonium bicarbonate followed by acetonitrile. 2) Reduced with 10mM dithiothreitol at 60°C followed by alkylation with 50mM iodoacetamide at RT. 3) Digested with trypsin (Promega, Madison, WI) at 37°C for 4h. 4) Quenched with formic acid and the supernatant was analyzed directly without further processing.

The digested protein samples were analyzed by nanoLC-MS/MS with a Waters NanoAcquity HPLC system interfaced to a ThermoFisher Q Exactive. Peptides were loaded on a trapping column and eluted over a 75µm analytical column at 350nL/min using a 2hr reverse phase gradient; both columns were packed with Luna C18 resin (Phenomenex, Torrance, CA). The mass spectrometer was operated in data-dependent mode, with the Orbitrap operating at 60,000 FWHM and 17,500 FWHM for MS and MS/MS respectively. The fifteen most abundant ions were selected for MS/MS. Data were searched using Mascot, and parsed into Scaffold (Proteome Software) for validation, filtering and to create a non-redundant list per sample. Data were filtered using 1% protein and peptide FDR and requiring at least two unique peptides per protein. A minimum of three spectral count values greater than 0 in at least one of the groups were considered significantly different, and a t-test was performed on these values. For protein pathway enrichment analysis, 222 proteins that were differentially expressed in AFSC-EVs were used as input for g:Profiler, and R package "ggplot2" was used to plot the top significantly enriched pathways for Biological Processes, Cellular Component, and Molecular Functions.

*RNA-sequencing of EV cargo*

Total RNA was isolated using the SeraMir Exosome RNA Purification Column kit (System Biosciences, Palo Alto, CA) according to the manufacturer's instructions. For each sample, 1µL of the final RNA eluate was used for measurement of small RNA concentration by Agilent Bioanalyzer Small RNA Assay using Bioanalyzer 2100 Expert instrument (Agilent Technologies, Santa Clara, CA).

Small RNA libraries were constructed with the CleanTag Small RNA Library Preparation Kit (TriLink, San Diego, CA) according to the manufacturer's protocol. The final purified library was quantified with High Sensitivity DNA Reagents (Agilent Technologies, Santa Clara, CA) and High Sensitivity DNA Chips (Agilent Technologies, Santa Clara, CA). The libraries were pooled, and the 140bp to 300bp region was size selected on an 8% TBE gel (Invitrogen, Carlsbad, CA). The size selected library is quantified with High Sensitivity DNA 1000 Screen Tape (Agilent Technologies, Carlsbad, CA), High Sensitivity D1000 reagents (Agilent

Technologies, Carlsbad, CA), and the TailorMix HT1 qPCR assay (SeqMatic, Fremont, CA), followed by a NextSeq High Output single-end sequencing run at SR75 using NextSeq 500/550 High Output v2 kit (Illumina, San Diego, CA) according to the manufacturer's instructions.

Following an initial quality assessment with FASTQC, n=3 biological replicates of AFSC-EVs and n=2 biological replicates of MSC-EVs were included in the final analysis. Bowtie2 was used to map the spike-in DNA, and the reads were trimmed and filtered to improve the quality of data input for read mapping using open-source tools (FastqMcf, cutadapt, PRINSEQ). After trimming, reads were mapped using open-source software (Bedtools/SAMtools). DESeq was used for differential expression analysis with default settings.

*Primary Lung Epithelial Cell RNA-sequencing Experiments*

Primary lung epithelial cells from nitrofen-exposed lungs and normal control lungs were isolated as described above, grown in PneumaCult™ Ex Plus Medium, and treated with 10% by volume AFSC-EVs or MSC-EVs. Total RNA extraction was conducted on n=6 biological replicates of each condition. RNA was extracted using Nucleospin miRNA kit (Macherey-Nagel, Düren, Germany) following supplier recommended protocols. Purified RNA was quantified using NanoDrop and Bioanalyzer was used to assess the quality of RNA. Samples with RNA integrity number of >9 were used to construct libraries. To construct RNA-seq libraries, we used an automated NEBNext Ultra II Directional with polyA isolation (New England BioLabs) using the Agilent NGS Workstation (Agilent Technologies) as per manufacturer's protocol. Briefly, 250 ng of total RNA spiked-in with SIRVs (Spike-in RNA Variant Control Mixes, Set3, Lexogen) as per manufacturer's protocol was used to generate cDNA. cDNA was amplified with 12 PCR cycles. The resulting libraries were quantified with Qubit DNA HS (ThermoFisher). Fragment sizes were analyzed on the Agilent Bioanalyzer using the High Sensitivity DNA assay prior to sequencing. Paired-end sequencing was performed by The Centre for Applied Genomics, The Hospital for Sick Children, Toronto, Canada on a NovaSeq 6000 S2 flowcell (Illumina) with a read length of 100 base pairs. 25~35 million paired end reads were obtained for each library. Sequencing quality was examined using FastQC and qualimap (see data file S2 for detailed QC metrics on sequencing libraries). Reads were aligned to the rat genome (rnor6, obtained from UCSC genome browser database) using a splice aware aligner, STAR (version 2.5.1b) with default settings. Reads were assigned to genes using featureCounts (version 1.5.3) with parameters "-p –B –s 2 –Q 255". Gene models were obtained from Ensembl (Rnor_6.0.91). Reads from ERCC (External RNA Controls Consortium, Ambion) spike-ins (included in SIRV Set3) were only used for QC purposes. For each sample, a linear model was fitted between log2 RPKM values and log2 expected RNA amount of the ERCC transcripts to evaluate the accuracy of the RNA-seq measurement. Number of ERCC transcripts detected (TPM > 0) and the corresponding $R^2$ values from the linear models are listed in data file S2. Normalized gene counts (reads per million mapped reads (RPKM) were calculated with R package "edgeR" (version 3.26.5) with "calcNormFactors" and "rpkm" functions. Only genes with RPKM > 1 in at least 12 samples were used for downstream analyses. All samples were used in the model fitting and dispersion estimation steps. The Quasi-likelihood F-test method was then used to identify the differentially expressed genes between pairs of conditions of interest (false discovery rate corrected (FDR) p-value < 0.1). Heatmaps were generated with R package "pheatmap with colors representing row-scaled RPKM values (Figure 4A). R package "fgsea' was used for GSEA analysis. For each pair-wise comparison, genes are ranked based on their fold changes (NA vs. N or NM vs. N). GMT files for "C2: curated gene sets" and "C5: GO gene sets" were obtained from the MSigDB collections and used separately in the analysis.

*Small RNA-sequencing on epithelial cells*

Small RNA isolated from n=4 matched samples of Nitrofen and Nitrofen+AFSC-EVs were subjected to miRNA-sequencing with the NEBNext small RNA kit. Sequencing libraries were quantified and size selected as described above, and sequenced on a single-end 50-bp rapid run flowcell. FastQC (v0.11.7) was used to examine the quality of approximately 1.4 million mapped reads per sample. BBDuk (BBMap suite v37.90) was used to trim adaptor sequences from reads with reference adapter sequences provided by BBMap suite and settings "hdist=1 mink=11" for small RNA-seq reads. For miRNA size specificity, only reads less than 23 nucleotides in length were retained. Following trimming, FastQC was used to examine the quality of trimmed sequenced reads. miRDeep2 (v2.0.0) mapper.pl was used with default parameters to map reads of at least 18 nucleotides in length to rat genome (rnor6). Known and novel miRNAs were identified using miRDeep2 main algorithm (miRDeep2.pl) with default parameters and known mature miRNAs for rat which were obtained from miRBase (v22.1). Only known and novel miRNAs with reported miRDeep score ≥ 2 were retained for downstream analysis.

*Counts processing and differential miRNA expression analysis*

Prior to differential miRNA expression analysis, read counts were scaled to sample library sizes and read counts per million (CPM) were calculated using R (v3.6.0) edgeR functions "cpm" and "calcNormFactors" (v3.26.5). Only miRNAs with CPM ≥ 1 in at least 3 samples within each condition were retained for downstream analysis. Differentially expressed miRNAs were identified using edgeR. Quasi-likelihood F-test method was used to test for differential expression (FDR < 0.1) for nitrofen-exposed and AFSC-EV exposed lung epithelial cells (NA) – nitrofen-exposed only (N) cells.

*miRNA-mRNA gene target correlation*

Rat orthologs of miRNA mRNA gene targets were determined from TargetScan (v7.2) and miRTarBase (release v8.0). Only TargetScan gene targets with weighted context score percentile $\geq 50$ were retained. Spearman's correlation was calculated using logCPM miRNA expression and logRPKM gene expression for a given pair. miRNA-gene pairs with Spearman's correlation coefficient (rho) $< 0$ and p-value $\leq 0.05$ were considered negatively correlated.

*Cargo-seq miRNA-mRNA interaction network*

miRNAs with a detected expression value $\geq 2$ in AFSC-EVs were considered in this analysis. miRNA-gene target pairs were determined as outlined above and shown as an interaction network generated using Cytoscape. Target genes that are differentially expressed (FDR $< 0.1$) and has lower expression in AFSC-EV-treated samples (log2FC $< 0$) are shown in the interaction network (blue nodes). miRNAs detected in AFSC-EVs with higher median logCPM expression in AFSC-EV-treated and nitrofen-treated primary cells compared to nitrofen-only treated cells are considered "miRNA up in primary cells" (green nodes). "miRNA in AFSC-EV cargo" (white nodes) represent miRNAs which are detected in AFSC-EV cargo and are not detected in AFSC-EV-treated epithelial cells or these miRNAs have lower median logCPM expression in AFSC-EV-treated and nitrofen-treated primary cells compared to nitrofen-only treated primary cells.

**Table S1**: Highlighted proteins expressed in AFSC-EVs and MSC-EVs.

| **Identified proteins** | **Molecular weight (kDa)** | **Average NSAF MSC** | **Average NSAF AFSC** | **p-value for fold change AFSC / MSC** |
|---|---|---|---|---|
| Anxa1 | 39 | 0.000 | 0.017 | 0.000 |
| Anxa11 | 54 | 0.000 | 0.001 | 0.122 |
| Anxa2 | 39 | 0.012 | 0.019 | 0.232 |
| Anxa3 | 36 | 0.000 | 0.001 | 0.374 |
| Anxa4 | 36 | 0.000 | 0.003 | 0.011 |
| Anxa5 | 36 | 0.000 | 0.012 | 0.000 |
| Anxa6 | 76 | 0.000 | 0.012 | 0.000 |
| Anxa7 | 50 | 0.000 | 0.002 | 0.000 |
| Cd63 | 26 | 0.000 | 0.001 | 0.001 |
| Celf1 | 52 | 0.000 | 0.000 | 0.374 |
| Hnrnpa1 | 34 | 0.000 | 0.001 | 0.157 |
| Hnrnpa2b1 | 32 | 0.000 | 0.001 | 0.118 |
| Hnrnpc | 33 | 0.000 | 0.000 | 0.374 |
| Hnrnpf | 46 | 0.000 | 0.001 | 0.001 |
| Hnrnph1 | 49 | 0.000 | 0.002 | 0.000 |
| Hnrnpk | 51 | 0.000 | 0.001 | 0.028 |
| Hnrnpl | 68 | 0.000 | 0.001 | 0.165 |
| Hnrnpm | 74 | 0.000 | 0.001 | 0.036 |
| Hnrnpu | 88 | 0.000 | 0.002 | 0.017 |
| Hnrnpul2 | 85 | 0.000 | 0.000 | 0.001 |
| Hspa1a | 70 | 0.000 | 0.003 | 0.000 |
| Hspa2 | 70 | 0.000 | 0.001 | 0.374 |
| Hspa4 | 94 | 0.000 | 0.000 | 0.374 |
| Hspa5 | 72 | 0.008 | 0.010 | 0.032 |
| Hspa8 | 71 | 0.011 | 0.012 | 0.575 |
| Hspa9 | 74 | 0.000 | 0.002 | 0.000 |

kDa: kilodaltons
NSAF: normalized spectral abundance factor
AFSCs: amniotic fluid stem cells
MSCs: mesenchymal stromal cells

**Table S2:** miRNAs related to lung development that are differentially expressed in AFSC-EVs over MSC-EVs.

| miRNA | miR ID | chr | start | end | type | MSCs | AFSCs | $log_2$Fold Change | adjusted p-value | Relevance |
|---|---|---|---|---|---|---|---|---|---|---|
| miR-17 | rno-miR-17-5p | chr15 | 103640915 | 103640937 | + | 59.87 | 22473.82 | 8.5522 | 0 | The mir 17-92 cluster and its paralogues:<br><br>- control FGF10-mediated embryonic lung epithelial branching morphogenesis (*69*);<br><br>- control lung progenitor cell proliferation and differentiation (*59, 69*);<br><br>- regulate embryonic growth, apoptosis, and fetal lung development (*9*);<br><br>- regulate surfactant protein C secretion (miR19b and 92a) (*70*);<br><br>- are involved in alveolarization processes (mir17) (*71*). |
| | rno-mir-17-2 | chrX | 140167738 | 140167815 | - | 59.87 | 22473.82 | 8.5522 | 0 | |
| | rno-mir-17-1 | chr15 | 103640902 | 103640985 | + | 65.28 | 22863.5 | 8.4522 | 0 | |
| | rno-miR-17-1-3p | chr15 | 103640952 | 103640973 | + | 0.71 | 177.19 | 7.958 | 0 | |
| miR-18 | rno-miR-18a-5p | chr15 | 103641054 | 103641076 | + | 3.07 | 953.84 | 8.2779 | 0 | |
| | rno-mir-18a | chr15 | 103641038 | 103641133 | + | 5.28 | 1034.37 | 7.6146 | 0 | |
| | rno-miR-18a-3p | chr15 | 103641095 | 103641115 | + | 2.2 | 81.58 | 5.2098 | 0 | |
| miR-19 | rno-miR-19a-5p | chr15 | 103641197 | 103641216 | + | 0 | 1.08 | Inf | 1 | |
| | rno-miR-19b-1-5p | chr15 | 103641502 | 103641523 | + | 0 | 1.05 | Inf | 1 | |
| | rno-mir-19a | chr15 | 103641185 | 103641266 | + | 2.71 | 1007.59 | 8.5407 | 0 | |
| | rno-miR-19a-3p | chr15 | 103641233 | 103641255 | + | 2.71 | 1006.52 | 8.5392 | 0 | |
| | rno-mir-19b-2 | chrX | 140167226 | 140167321 | - | 16.7 | 5235.41 | 8.2922 | 0 | |
| | rno-miR-19b-3p | chr15 | 103641540 | 103641562 | + | 16.7 | 5235.41 | 8.2922 | 0 | |
| | rno-mir-19b-1 | chr15 | 103641487 | 103641573 | + | 16.7 | 5231.7 | 8.2912 | 0 | |
| miR-20 | rno-miR-20a-3p | chr15 | 103641408 | 103641428 | + | 0 | 1.6 | Inf | 0.8997 | |
| | rno-miR-20a-5p | chr15 | 103641372 | 103641394 | + | 30.04 | 19844.96 | 9.3677 | 0 | |
| | rno-mir-20a | chr15 | 103641357 | 103641441 | + | 30.04 | 19844.96 | 9.3677 | 0 | |
| | rno-miR-20b-5p | chrX | 140167407 | 140167429 | - | 0.51 | 21.61 | 5.3986 | 0.0038 | |
| | rno-mir-20b | chrX | 140167365 | 140167436 | - | 0.51 | 21.61 | 5.3986 | 0.0038 | |
| miR-92 | rno-miR-92b-5p | chr2 | 207955729 | 207955752 | - | 0.2 | 1.62 | 3.0149 | 1 | |
| | rno-miR-92a-1-5p | chr15 | 103641619 | 103641641 | + | 2.2 | 15.58 | 2.8214 | 0.1041 | |
| | rno-mir-92a-1 | chr15 | 103641609 | 103641686 | + | 1466.75 | 3152.85 | 1.104 | 0.0346 | |
| | rno-miR-92a-3p | chr15 | 103641656 | 103641676 | + | 447.53 | 519.1 | 0.214 | 1 | |
| | rno-mir-92a-2 | chrX | 140167096 | 140167187 | - | 447.53 | 519.1 | 0.214 | 1 | |
| | rno-mir-92b | chr2 | 207955680 | 207955762 | - | 404.68 | 109.76 | -1.8824 | 0.1754 | |
| | rno-miR-92b-3p | chr2 | 207955690 | 207955711 | - | 404.48 | 108.14 | -1.9032 | 0.1696 | |
| miR-106 | rno-miR-106b-5p | chr12 | 21365432 | 21365452 | - | 12.55 | 2451.88 | 7.6103 | 0 | |
| | rno-mir-106b | chr12 | 21365382 | 21365463 | - | 302.19 | 12390.58 | 5.3576 | 0 | |
| | rno-miR-106b-3p | chr12 | 21365391 | 21365412 | - | 289.64 | 9939.25 | 5.1008 | 0 | |
| miR-363 | rno-miR-363-3p | chrX | 140166956 | 140166976 | - | 3.21 | 3.72 | 0.2128 | 1 | |
| | rno-mir-363 | chrX | 140166945 | 140167031 | - | 4.49 | 3.72 | -0.2719 | 1 | |

| | | | | | | | | | |
|---|---|---|---|---|---|---|---|---|---|
| miR-93 | rno-miR-93-5p | chr12 | 21365223 | 21365245 | - | 953.63 | 108642.09 | 6.8319 | 0 |
| | rno-mir-93 | chr12 | 21365173 | 21365259 | - | 955.42 | 108717.23 | 6.8302 | 0 |
| | rno-miR-93-3p | chr12 | 21365185 | 21365207 | - | 1.79 | 75.15 | 5.3896 | 0 |
| miR-25 | rno-miR-25-3p | chr12 | 21364981 | 21365002 | - | 5238.63 | 83820.36 | 4 | 0 |
| | rno-mir-25 | chr12 | 21364970 | 21365053 | - | 5262.06 | 83911.87 | 3.9952 | 0 |
| | rno-miR-25-5p | chr12 | 21365019 | 21365040 | - | 23.43 | 90.98 | 1.9574 | 0.0033 |

| | | | | | | | | | | |
|---|---|---|---|---|---|---|---|---|---|---|
| miR-7 | rno-miR-7b | chr9 | 9803964 | 9803986 | - | 0 | 66.56 | Inf | 0 | - miRNAs involved in surfactant protein C secretion and expressed in human ATI and/or ATII cells (*70*). |
| | rno-mir-7b | chr9 | 9803905 | 9804014 | - | 0 | 66.56 | Inf | 0 | |
| | rno-miR-7a-2-3p | chr1 | 141551398 | 141551419 | + | 0 | 2.68 | Inf | 0.6528 | |
| | rno-mir-7a-2 | chr1 | 141551342 | 141551436 | + | 111.49 | 133696.6 | 10.2279 | 0 | |
| | rno-miR-7a-5p | chr1 | 141551360 | 141551382 | + | 119.28 | 133897.01 | 10.1325 | 0 | |
| | rno-mir-7a-1 | chr17 | 8879389 | 8879485 | + | 125.25 | 133996 | 10.0632 | 0 | |
| | rno-miR-7a-1-3p | chr17 | 8879448 | 8879469 | + | 5.97 | 97.89 | 4.0358 | 0 | |
| let7-f-1 and let7-f-2 | rno-let-7f-1-3p | chr17 | 18474397 | 18474417 | + | 0 | 37.3 | Inf | 4.00E-04 | |
| | rno-let-7f-1 | chr17 | 18474334 | 18474422 | + | 6660.85 | 188375.58 | 4.8218 | 0 | |
| | rno-let-7f-5p | chr17 | 18474341 | 18474362 | + | 7068.36 | 189106.28 | 4.7417 | 0 | |
| | rno-let-7f-2 | chrX | 21868509 | 21868591 | - | 7072.71 | 189158.31 | 4.7412 | 0 | |
| | rno-let-7f-2-3p | chrX | 21868514 | 21868534 | - | 3.59 | 42.95 | 3.5827 | 0 | |
| rno-mir-219a-1 | rno-mir-219a-1 | chr20 | 5895489 | 5895598 | - | 10.69 | 85.29 | 2.9963 | 0 | |
| | rno-miR-219a-1-3p | chr20 | 5895516 | 5895537 | - | 10.69 | 83.16 | 2.9598 | 0 | |
| miR-103 | rno-mir-103-2 | chr3 | 130329252 | 130329337 | + | 3229.37 | 40889.34 | 3.6624 | 0 | |
| | rno-miR-103-3p | chr10 | 20480402 | 20480424 | + | 3229.37 | 40883.42 | 3.6622 | 0 | |
| | rno-mir-103-1 | chr10 | 20480351 | 20480436 | + | 3234.27 | 40892.4 | 3.6603 | 0 | |
| miR-125b-1 | rno-miR-125b-1-3p | chr8 | 44272900 | 44272921 | + | 7472.2 | 19346.02 | 1.3724 | 0.002 | |

| | | | | | | | | | | |
|---|---|---|---|---|---|---|---|---|---|---|
| | rno-mir-125b-1 | chr8 | 44272846 | 44272932 | + | 7718.59 | 19862 | 1.3636 | 0.0022 | |
| mir-129-1 | rno-miR-129-1-3p | chr4 | 56049143 | 56049161 | + | 0 | 2.66 | Inf | 0.5709 | |
| | rno-mir-129-1 | chr4 | 56049095 | 56049166 | + | 369.44 | 5715.96 | 3.9516 | 0 | |
| miR-542 | rno-miR-542-3p | chrX | 152777501 | 152777522 | + | 47.45 | 14042.5 | 8.2091 | 0 | |
| | rno-mir-542-1 | chrX | 152777453 | 152777531 | + | 60.06 | 14313.6 | 7.8968 | 0 | |
| | rno-mir-542-2 | chrX | 152784192 | 152784270 | + | 60.06 | 14313.6 | 7.8968 | 0 | |
| | rno-mir-542-3 | chrX | 153215172 | 153215250 | + | 60.06 | 14313.6 | 7.8968 | 0 | |
| | rno-miR-542-5p | chrX | 152777463 | 152777484 | + | 12.6 | 265.18 | 4.395 | 0 | |
| miR-592 | rno-mir-592 | chr4 | 54925353 | 54925448 | - | 18.39 | 4706.09 | 7.9992 | 0 | |
| | rno-miR-592 | chr4 | 54925409 | 54925431 | - | 18.39 | 4705.01 | 7.9989 | 0 | |
| miR-138 | rno-miR-138-5p | chr19 | 11127530 | 11127552 | - | 5.59 | 109.53 | 4.2925 | 0 | - miRNAs that regulate late stage murine lung development and are significantly different with sex and gestational age in E15-E18 lungs (*72*). |
| | rno-mir-138-1 | chr8 | 130886768 | 130886866 | + | 5.85 | 109.53 | 4.2279 | 0 | |
| | rno-mir-138-2 | chr19 | 11127479 | 11127560 | - | 6.87 | 124.48 | 4.1794 | 0 | |
| | rno-miR-138-2-3p | chr19 | 11127485 | 11127505 | - | 1.28 | 19.31 | 3.9144 | 0.0039 | |
| miR-182 | rno-miR-182 | chr4 | 57221577 | 57221601 | - | 40.34 | 994.77 | 4.6241 | 0 | |
| | rno-mir-182 | chr4 | 57221574 | 57221635 | - | 40.34 | 994.77 | 4.6241 | 0 | |
| mir-296 | rno-miR-296-5p | chr3 | 178408833 | 178408853 | - | 0.6 | 41.26 | 6.1007 | 0 | |
| | rno-mir-296 | chr3 | 178408788 | 178408865 | - | 116.43 | 516.99 | 2.1507 | 0 | |
| | rno-miR-296-3p | chr3 | 178408798 | 178408819 | - | 115.83 | 471.97 | 2.0267 | 4.00E-04 | |
| miR-471 | rno-miR-471-3p | chrX | 151267019 | 151267038 | - | 0 | 66.6 | Inf | 0 | - miRNAs involved in alveolarization and significantly changed in P10 or P20 lungs in a model of intrauterine growth restriction (*71*). |
| | rno-mir-471 | chrX | 151267006 | 151267083 | - | 0.71 | 691.21 | 9.9218 | 0 | |
| | rno-miR-471-5p | chrX | 151267052 | 151267073 | - | 0.71 | 621.36 | 9.7681 | 0 | |
| miR-455 | rno-miR-455-5p | chr5 | 83211466 | 83211487 | + | 120.43 | 1324.53 | 3.4593 | 0 | |
| miR-322 | rno-miR-322-3p | chrX | 152773528 | 152773547 | + | 62.07 | 6528.84 | 6.7167 | 0 | |
| | rno-miR-322-5p | chrX | 152773490 | 152773511 | + | 11.56 | 2847.63 | 7.9448 | 0 | |
| | rno-mir-322-1 | chrX | 152773468 | 152773562 | + | 73.63 | 9392 | 6.995 | 0 | |
| | rno-mir-322-2 | chrX | 153211185 | 153211279 | + | 73.63 | 9392 | 6.995 | 0 | |
| | rno-miR-322-3p | chrX | 152773528 | 152773547 | + | 62.07 | 6528.84 | 6.7167 | 0 | |
| miR-183 | rno-mir-183 | chr4 | 57225370 | 57225479 | - | 13.24 | 316.45 | 4.5792 | 0 | |
| | rno-miR-183-5p | chr4 | 57225432 | 57225453 | - | 13.24 | 315.9 | 4.5767 | 0 | |
| miR-214 | rno-miR-214-5p | chr13 | 85024828 | 85024844 | + | 50.79 | 1468.47 | 4.8535 | 0 | |
| miR-130 | rno-miR-130a-5p | chr3 | 78661749 | 78661770 | - | 0 | 10.72 | Inf | 0.0051 | |

| | | | | | | | | | | |
|---|---|---|---|---|---|---|---|---|---|---|
| | rno-miR-130b-3p | chr11 | 91182370 | 91182391 | + | 251.24 | 18538.82 | 6.2053 | 0 | |
| | rno-mir-130b | chr11 | 91182320 | 91182401 | + | 254.57 | 18584.85 | 6.1899 | 0 | |
| | rno-mir-130a | chr3 | 78661697 | 78661784 | - | 1092.48 | 20003.91 | 4.1946 | 0 | |
| | rno-miR-130a-3p | chr3 | 78661709 | 78661730 | - | 1092.48 | 19991.05 | 4.1937 | 0 | |
| | rno-miR-130b-5p | chr11 | 91182332 | 91182353 | + | 3.33 | 44.39 | 3.7371 | 0.09 | |
| miR-463 | rno-miR-463-3p | chrX | 151270941 | 151270962 | - | 0 | 2165.36 | Inf | 0 | |
| | rno-mir-463 | chrX | 151270931 | 151271008 | - | 0.26 | 3748.68 | 13.8374 | 0 | |
| | rno-miR-463-5p | chrX | 151270979 | 151270999 | - | 0.26 | 1582.78 | 12.5935 | 0 | |
| miR-465 | rno-mir-465 | chrX | 151292178 | 151292254 | - | 0 | 2167.1 | Inf | 0 | |
| | rno-miR-465-5p | chrX | 151292223 | 151292244 | - | 0 | 2096.67 | Inf | 0 | |
| | rno-miR-465-3p | chrX | 151292189 | 151292210 | - | 0 | 68.81 | Inf | 0 | |
| miR-471 | rno-miR-471-3p | chrX | 151267019 | 151267038 | - | 0 | 66.6 | Inf | 0 | |
| | rno-mir-471 | chrX | 151267006 | 151267083 | - | 0.71 | 691.21 | 9.9218 | 0 | |
| | rno-miR-471-5p | chrX | 151267052 | 151267073 | - | 0.71 | 621.36 | 9.7681 | 0 | |
| miR-741 | rno-miR-741-3p | chrX | 151269275 | 151269296 | - | 0 | 816.46 | Inf | 0 | |
| | rno-mir-741 | chrX | 151269258 | 151269353 | - | 0 | 816.46 | Inf | 0 | - miRNAs involved in the regulation of pluripotency and the reprogramming process in rats (*73*). |
| miR-743b | rno-mir-743b | chrX | 151248558 | 151248634 | - | 0 | 3809.35 | Inf | 0 | |
| | rno-miR-743b-5p | chrX | 151248603 | 151248624 | - | 0 | 2753.09 | Inf | 0 | |
| | rno-miR-743b-3p | chrX | 151248568 | 151248589 | - | 0 | 1054.16 | Inf | 0 | |
| miR-871 | rno-mir-871 | chrX | 151282689 | 151282765 | - | 0 | 7147.82 | Inf | 0 | |
| | rno-miR-871-5p | chrX | 151282732 | 151282755 | - | 0 | 7109.67 | Inf | 0 | |
| | rno-miR-871-3p | chrX | 151282700 | 151282721 | - | 0 | 38.15 | Inf | 0 | |
| miR-881 | rno-mir-881 | chrX | 151278812 | 151278888 | - | 0 | 2210.79 | Inf | 0 | |
| | rno-miR-881-3p | chrX | 151278823 | 151278844 | - | 0 | 1995.88 | Inf | 0 | |
| | rno-miR-881-5p | chrX | 151278860 | 151278878 | - | 0 | 214.9 | Inf | 0 | |
| miR-883 | rno-mir-883 | chrX | 151255943 | 151256019 | - | 0 | 185.76 | Inf | 0 | |
| | rno-miR-883-3p | chrX | 151255953 | 151255974 | - | 0 | 175.6 | Inf | 0 | |
| | rno-miR-883-5p | chrX | 151255988 | 151256010 | - | 0 | 10.16 | Inf | 0.0072 | |
| miR-3580 | rno-mir-3580 | chrX | 151290426 | 151290506 | - | 0 | 4306.44 | Inf | 0 | |
| | rno-miR-3580-3p | chrX | 151290438 | 151290459 | - | 0 | 4168.11 | Inf | 0 | |
| | rno-miR-3580-5p | chrX | 151290472 | 151290493 | - | 0 | 138.34 | Inf | 0 | |

miR: miRNA; AFSCs: amniotic fluid stem cells; MSCs: mesenchymal stem cells; chr: chromosome; inf: infinity; ATI: alveolar type I cells; ATII: alveolar type II cells.

**Table S3:** miRNAs known to be involved in pulmonary hypoplasia and present in AFSC-EVs.

| miRNA | miR ID | chr | start | end | type | MSCs | AFSCs | $log_2$Fold Change | adjusted p-value | Relevance |
|---|---|---|---|---|---|---|---|---|---|---|
| miR-200 | rno-miR-200a-5p | chr5 | 176963441 | 176963461 | - | 0.51 | 0 | Inf | 1 | - miRNAs dysregulated in pulmonary hypoplasia secondary to CDH (*12, 74-76*). |
| | rno-mir-200b | chr5 | 176964166 | 176964260 | - | 4.65 | 0 | Inf | 0.4715 | |
| | rno-miR-200b-3p | chr5 | 176964182 | 176964204 | - | 1.54 | 0 | Inf | 1 | |
| | rno-miR-200b-5p | chr5 | 176964219 | 176964240 | - | 3.12 | 0 | Inf | 0.7365 | |
| | rno-miR-200c-5p | chr4 | 224254426 | 224254446 | - | 0 | 0.55 | Inf | 1 | |
| | rno-mir-200c | chr4 | 224254382 | 224254450 | - | 12.7 | 65.95 | 2.376 | 0.0014 | |
| | rno-miR-200c-3p | chr4 | 224254386 | 224254406 | - | 12.7 | 65.95 | 2.376 | 0.0014 | |
| | rno-miR-200a-3p | chr5 | 176963402 | 176963423 | - | 5.57 | 3.8 | -0.5508 | 1 | |
| | rno-mir-200a | chr5 | 176963388 | 176963476 | - | 6.08 | 3.8 | -0.6778 | 1 | |
| miR-10 | rno-miR-10b-3p | chr3 | 68113726 | 68113746 | + | 27.86 | 172.28 | 2.6286 | 1.00E-04 | |
| | rno-mir-10b | chr3 | 68113662 | 68113770 | + | 29928.19 | 88517.54 | 1.5645 | 4.00E-04 | |
| | rno-miR-10b-5p | chr3 | 68113689 | 68113710 | + | 29900.33 | 88350.61 | 1.5631 | 4.00E-04 | |
| | rno-miR-10a-3p | chr10 | 83968744 | 83968765 | + | 4.78 | 2.66 | -0.8426 | 1 | |
| | rno-mir-10a | chr10 | 83968682 | 83968791 | + | 727.64 | 88.91 | -3.0328 | 0.0256 | |
| | rno-miR-10a-5p | chr10 | 83968703 | 83968725 | + | 722.86 | 86.25 | -3.0671 | 0.0232 | |
| miR-33 | rno-miR-33-3p | chr7 | 123415348 | 123415368 | + | 0 | 4.83 | Inf | 0.1942 | - miRNAs downregulated in hypoplastic lungs of experimental CDH (nitrofen model) (*10*). |
| | rno-mir-33 | chr7 | 123415303 | 123415371 | + | 3.94 | 49.49 | 3.6504 | 0 | |
| | rno-miR-33-5p | chr7 | 123415308 | 123415328 | + | 2.43 | 33.88 | 3.8032 | 2.00E-04 | |
| miR-193 | rno-mir-193b | chr14 | 30144277 | 30144359 | - | 43.73 | 552.81 | 3.66 | 0 | |
| | rno-miR-193b-3p | chr14 | 30144291 | 30144309 | - | 2.88 | 543.63 | 7.5585 | 0 | |
| | rno-miR-193a-3p | chr10 | 64583983 | 64584004 | - | 2.32 | 22.62 | 3.2878 | 0.0117 | |
| | rno-mir-193a | chr10 | 64583972 | 64584057 | - | 60.8 | 160.84 | 1.4034 | 0.0185 | |
| | rno-miR-193a-5p | chr10 | 64584017 | 64584038 | - | 56.55 | 138.22 | 1.2894 | 0.0417 | |
| | rno-miR-193b-5p | chr14 | 30144328 | 30144345 | - | 40.85 | 6.45 | -2.6624 | 0.1982 | |
| miR-338 | rno-mir-338 | chr10 | 108793410 | 108793475 | - | 0 | 16.14 | Inf | 2.00E-04 | |
| | rno-miR-338-3p | chr10 | 108793413 | 108793435 | - | 0 | 8.1 | Inf | 0.0761 | |
| | rno-miR-338-5p | chr10 | 108793449 | 108793470 | - | 0 | 8.04 | Inf | 0.0266 | |
| miR-30a | rno-mir-30a | chr9 | 28377823 | 28377893 | + | 6702.06 | 91515.55 | 3.7713 | 0 | |
| | rno-miR-30a-5p | chr9 | 28377828 | 28377849 | + | 6239.92 | 88950.46 | 3.8334 | 0 | |
| | rno-miR-30a-3p | chr9 | 28377869 | 28377890 | + | 462.13 | 2565.09 | 2.4726 | 0 | |

| | | | | | | | | | |
|---|---|---|---|---|---|---|---|---|---|
| and | | | | | | | | | |
| miR-30c | rno-mir-30c-2 | chr9 | 28397564 | 28397647 | + | 126.18 | 1532.57 | 3.6024 | 0 |
| | rno-mir-30c-1 | chr5 | 143494757 | 143494845 | - | 113.57 | 1523.39 | 3.7457 | 0 |
| | rno-miR-30c-5p | chr5 | 143494807 | 143494829 | - | 110.28 | 1483.2 | 3.7494 | 0 |
| | rno-miR-30c-1-3p | chr5 | 143494769 | 143494790 | - | 3.28 | 40.71 | 3.6316 | 1.00E-04 |
| | rno-miR-30c-2-3p | chr9 | 28397617 | 28397638 | + | 15.9 | 49.37 | 1.6346 | 0.0871 |
| miR-22 | rno-miR-22-3p | chr10 | 62013698 | 62013719 | + | 11519.53 | 80600.86 | 2.8067 | 0 |
| | rno-miR-22-5p | chr10 | 62013660 | 62013681 | + | 43.31 | 499.36 | 3.5272 | 0 |
| | rno-mir-22 | chr10 | 62013642 | 62013736 | + | 11562.84 | 81100.22 | 2.8102 | 0 |
| miR-532 | rno-mir-532 | chrX | 16894994 | 16895072 | + | 9.75 | 128.96 | 3.7249 | 0 |
| | rno-miR-532-5p | chrX | 16895004 | 16895025 | + | 3.92 | 94.01 | 4.5842 | 0 |
| | rno-miR-532-3p | chrX | 16895041 | 16895062 | + | 5.83 | 34.95 | 2.5827 | 0.013 |
| miR-28 | rno-miR-28-3p | chr11 | 81345228 | 81345249 | + | 1115.34 | 34443.35 | 4.9487 | 0 |
| | rno-miR-28-5p | chr11 | 81345188 | 81345209 | + | 76.11 | 3651.38 | 5.5842 | 0 |
| | rno-mir-28 | chr11 | 81345175 | 81345260 | + | 1191.45 | 38095.82 | 4.9988 | 0 |
| miR-362 | rno-mir-362 | chrX | 16902294 | 16902358 | + | 0 | 115.82 | Inf | 0 |
| | rno-miR-362-3p | chrX | 16902335 | 16902356 | + | 0 | 114.72 | Inf | 0 |
| | rno-miR-362-5p | chrX | 16902298 | 16902321 | + | 0 | 1.09 | Inf | 1 |
| miR-3559 | rno-mir-3559 | chrX | 76261608 | 76261722 | - | 26.31 | 1132.64 | 5.4279 | 0 |
| | rno-miR-3559-3p | chrX | 76261633 | 76261654 | - | 21.44 | 712.18 | 5.0536 | 0 |
| | rno-miR-3559-5p | chrX | 76261673 | 76261694 | - | 4.87 | 420.45 | 6.4331 | 0 |

miR: miRNA; AFSCs: amniotic fluid stem cells; MSCs: mesenchymal stem cells; chr: chromosome; inf: infinity.

**Table S4:** Genes differentially expressed in nitrofen-exposed lung epithelial cells

| Gene | Name | Link to lung development or epithelial homeostasis | Ref. |
|---|---|---|---|
| **Down-regulated** | | | |
| Nkx2.1 | NK2 Homeobox 1 | Essential regulator of lung development and a marker of early lung epithelial progenitor | (*77*) |
| Hipk2 | Homeodomain Interacting Protein Kinase 2 | NKX2-1 binding partner | (*78*) |
| Alcam | Activated Leukocyte Cell Adhesion Molecule | Marker of type II alveolar epithelial cells | (*79*) |
| Mxd4 | MAX Dimerization Protein 4 | Highly prioritized gene in neonates with pulmonary hypoplasia /CDH | (*80*) |
| Arg2 | Arginase 2 | Overexpressed gene in neonates with pulmonary hypoplasia/CDH | (*81*) |
| Fgfr3 | Fibroblast Growth Factor Receptor 3 | Upregulation of FGFR3 disrupts alveologenesis in experimental and human pulmonary hypoplasia/CDH. | (*80*, *82*) |
| Sult1a1 | Sulfotransferase Family 1A Member 1 | Upregulated in hypoxic lung injury, and required for nitrofen activation and mutagenicity | (*83, 84*) |
| Dusp1 | Dual Specificity Phosphatase 1 | Negatively regulates autophagy, a process required for fetal lung branching morphogenesis | (*85, 86*) |
| Sqstm1 | Sequestosome 1 | Marker of autophagy impairment increased in stressed conditions | (*86, 87*) |
| Ncao4 | Nuclear Receptor Coactivator 4 | Involved in ferritin turnover and autophagy regulation | (*88*) |
| Herpud1 | Homocysteine Inducible ER Protein With Ubiquitin Like Domain 1 | Involved in ER stress response, a critical process for tissue homeostasis that is dysregulated in pulmonary hypoplasia | (*40, 89*) |
| Cdkn1c | Cyclin Dependent Kinase | Highly expressed in pulmonary hypoplasia | (*90*) |

| | | | |
|---|---|---|---|
| | Inhibitor 1C | | |
| Fuca2 | Alpha-L-Fucosidase 2 | Involved in repair of damaged lung epithelial cells | (*91*) |
| Clic3 | Chloride Intracellular Channel 3 | Activator of MAPK signaling, a pathway that is upregulated in nitrofen-exposed lung epithelial cells | (*61, 92*) |
| Sesn1 | Sestrin 1 | Repressor of PDGFRβ signaling, a pathway that is dysregulated in nitrofen lungs | (*93*) |
| Flcn | Folliculin | Involved in EGFR signaling, a pathway that is down-regulated in pulmonary hypoplasia. | (*94, 95*) |
| **Up-regulated** | | | |
| Flna | Filamin A | Actin-binding protein involved in ciliogenesis, FLNA mutations are associated with severe diffuse lung disease and alveolar simplification | (*96, 97*) |
| Pdlim5 | PDZ And LIM Domain 5 | Scaffolding protein that is required for TGF-β/Smad signaling in alveolar epithelial cells, and that prevents hypoxia-induced pulmonary hypertension | (*98*) |
| Gldn | Gliomedin | ECM matrix glycoprotein (up-regulated by TGF-β signaling) | (*99*) |
| Igf1r | Insulin Like Growth Factor 1 Receptor | Involved in epithelial proliferation and differentiation, (downregulated in nitrofen) | (*32*) |
| Gnaq | G Protein Subunit Alpha Q | Nucleotide-binding protein expressed in alveolar Type II epithelial cells that controls surfactant homeostasis | (*100*) |
| Myh9l1 | Myosin Heavy Chain 9 non-muscle-like 1 | Protein important for epithelial cell tight junction formation | (*101*) |
| Clic1 | Chloride Intracellular Channel 1 | Mitochondria and ER chloride channel protein that regulates redox balance and ER stress | (*40, 102*) |
| Hikeshi | Heat Shock Protein | Nuclear transport receptor required for organization and | (*103*) |

| | | | |
|---|---|---|---|
| | Nuclear Import Factor Hikeshi | function of the secretory apparatus in club cells | |
| Cxcl2 | C-X-C Motif Chemokine Ligand 2 | Cytokine that stimulates proliferation in rat alveolar epithelial cells. | (*104*) |

**Table S5: Inter-species conservation of top enriched miRNA in human AFSC-EVs**

| Human miRNA | Rabbit miRNA orthologue | Sequence |
|---|---|---|
| hsa-let-7b-5p | ocu-let-7b | 5`-UGAGGUAGUAGGUUGUGUGGUU-3` |
| hsa-miR-23a-3p | ocu-miR-23b-3p | 5`-AUCACAUUGCCAGGGAUUUCC-3` |
| hsa-miR-24-3p | ocu-miR-24-3p | 5`-UGGCUCAGUUCAGCAGGAACAG-3` |
| hsa-miR-31-5p | ocu-miR-31-5p | 5`-AGGCAAGAUGCUGGCAUAGCU-3` |
| hsa-miR-92a-3p | ocu-miR-92a-3p | 5`-UAUUGCACUUGUCCCGGCCUGU-3` |
| hsa-miR-100-5p | ocu-miR-100-5p | 5`-AACCCGUAGAUCCGAACUUGUG-3` |
| hsa-miR-103a-3p | ocu-miR-103a-3p | 5`-AGCAGCAUUGUACAGGGCUAUGA-3` |
| hsa-miR-107 | ocu-miR-107-3p | 5`-AGCAGCAUUGUACAGGGCUAUCA-3` |
| hsa-miR-221-3p | ocu-miR-221-3p | 5`-AGCUACAUUGUCUGCUGGGUUUC-3` |
| hsa-miR-222-3p | ocu-miR-222-3p | 5`-AGCUACAUCUGGCUACUGGGU-3` |
| hsa-miR-23b-3p | ocu-miR-23b-3p | 5`-AUCACAUUGCCAGGGAUUACCAC-3` |
| hsa-miR-125b-5p | ocu-miR-125b-5p | 5`-UCCCUGAGACCCUAACUUGUGA-3` |
| hsa-miR-145-5p | ocu-miR-145-5p | 5`-GUCCAGUUUUCCCAGGAAUCCCU-3` |
| hsa-miR-320c | ocu-miR-197-3p | 5`-AAAAGCUGGGUUGAGAGGGU-3` |
| hsa-miR-3613-3p | ocu-miR-3613-3p | 5`-ACAAAAAAAAAAGCCCAACCCUUC-3` |

**Table S6: Primer sequences used in this study**

| Target | Primer sequence |
|---|---|
| rat Gapdh-F | 5`-GGGTGTGAACCACGAGAAAT-3` |
| rat Gapdh-R | 5`-ACTGTGGTCATGAGCCCTTC-3` |
| rat Fgf10-F | 5`-CCACATACATTTGCCTGCCG-3` |
| rat Fgf10-R | 5`-GGGGAAACTCTATGGCTCAAAAG-3` |
| rat Vegfa-F | 5`-AGAAAGCCCATGAAGTGGTGA-3` |
| rat Vegfa -R | 5`-TCTCATCGGGGTACTCCTGG-3` |
| rat Flt1-F | 5`-GTACCTCACCGTGCAAGGAA-3` |
| rat Flt1-R | 5`-TTCGGAAGAAGACCGCTTCA-3` |
| rat Kdr-F | 5`-CTGCAGGACCAAGGCAACTA-3` |
| rat Kdr-R | 5`-CATGCGCTCTAGGATGACGA-3` |
| rabbit RPLP0-F | 5`-CTGTGCCAGCTCAGAACACT-3` |
| rabbit RPLP0-R | 5`-TGCACGTCGCTCAGGATTTC-3` |
| rabbit PLIN-2-F | 5`-TGCTGAGCACATCGAGTCAC-3` |
| rabbit PLIN-2-R | 5`-ATGTTGGACAGGAGGCTGTG-3` |
| rabbit BMP2-F | 5`-GGAAGCTTTGGGAGACGACA-3` |
| rabbit BMP2-R | 5`-TTTCGAGTTGGCTGTTGCAG-3` |
| rabbit BMP4-F | 5`-CTTCCACCACGAAGAACATCTG-3` |
| rabbit BMP4-R | 5`-ATGGCCTCGTTCTCTGGGAT-3` |
| rabbit Id1-F | 5`-TTCTACAACCGTCTCCTGCG-3` |
| rabbit Id1-R | 5`-CTGGCGACCTTCATGGTTCT-3` |

**Table S7: Details of antibodies used in this study**

| Target | Antibody | Company | Lung explants | | Organoids | | Primary epithelial cells | |
|---|---|---|---|---|---|---|---|---|
| | | | 1° | 2° | 1° | 2° | 1° | 2° |
| SPC | ab40879 | Abcam (Cambridge, UK) | 1:500 | 1:1,000 | 1:200 | 1:1,000 | 1:200 | 1:1,000 |
| Sox9 | HPA001758 | SigmaAldrich (St Louis, MO) | 1:500 | 1:1,000 | - | - | - | - |
| Ki67 | ab15580 | Abcam (Cambridge, UK) | - | - | 1:100 | 1:1,000 | - | - |
| CC10 | sc-365992 | SantaCruz Biotechnology (Dallas, TX) | - | - | 1:50 | 1:1,000 | - | - |
| Vimentin | ab92547 | Abcam (Cambridge, UK) | - | - | - | - | 1:100 | 1:1,000 |

| Target | Antibody | Company | Western Blot | | ImmunoEM | |
|---|---|---|---|---|---|---|
| | | | 1° | 2° | 1° | 2° |
| TSG101 | sc-7964 | Santa Cruz Biotechnology, Dallas, TX | 1:500 | 1:3,000 | 1:100 | - |
| Goat-anti-mouse IgG (H&L) | 25128 EM-grade 10nm gold tag | Electron Microscopy Sciences, Hatfield, PA | - | - | - | 1:25 |
| RNase | PA578151 | Thermofisher Scientific, Waltham, Massachusetts | - | - | 1:100 | - |
| Goat-anti-rabbit IgG (H&L) | 25116 EM-grade 25nm gold tag | Electron Microscopy Sciences, Hatfield, PA | - | - | - | 1:25 |
| CD63 | EXOAB-KIT-1 | System Biosciences, Palo Alto, CA | 1:1,000 | 1:10,000 | - | - |
| Flo-1 | 610820 | BD Transduction Laboratories, San Jose, CA | 1:1,000 | 1:3,000 | - | - |
| Hsp70 | EXOAB-KIT-1 | System Biosciences, Palo Alto, CA | 1:1,000 | 1:10,000 | - | - |

SPC: surfactant protein C,
Sox9: SRY-Box 9,
Ki67: marker of proliferation Ki67,
CC10: Clara Cells 10 KDa Secretory Protein,
TSG101: Tumor susceptibility gene 101,
Flo-1: Flotillin 1,

Hsp70: Heat Shock Protein 70
