## Supplementary figures and images for "Impaired Fetal Lung Development can be Rescued by Administration of Extracellular Vesicles Derived from Amniotic Fluid Stem Cells"

### bioRxiv-supp_Sup Fig 1.tif

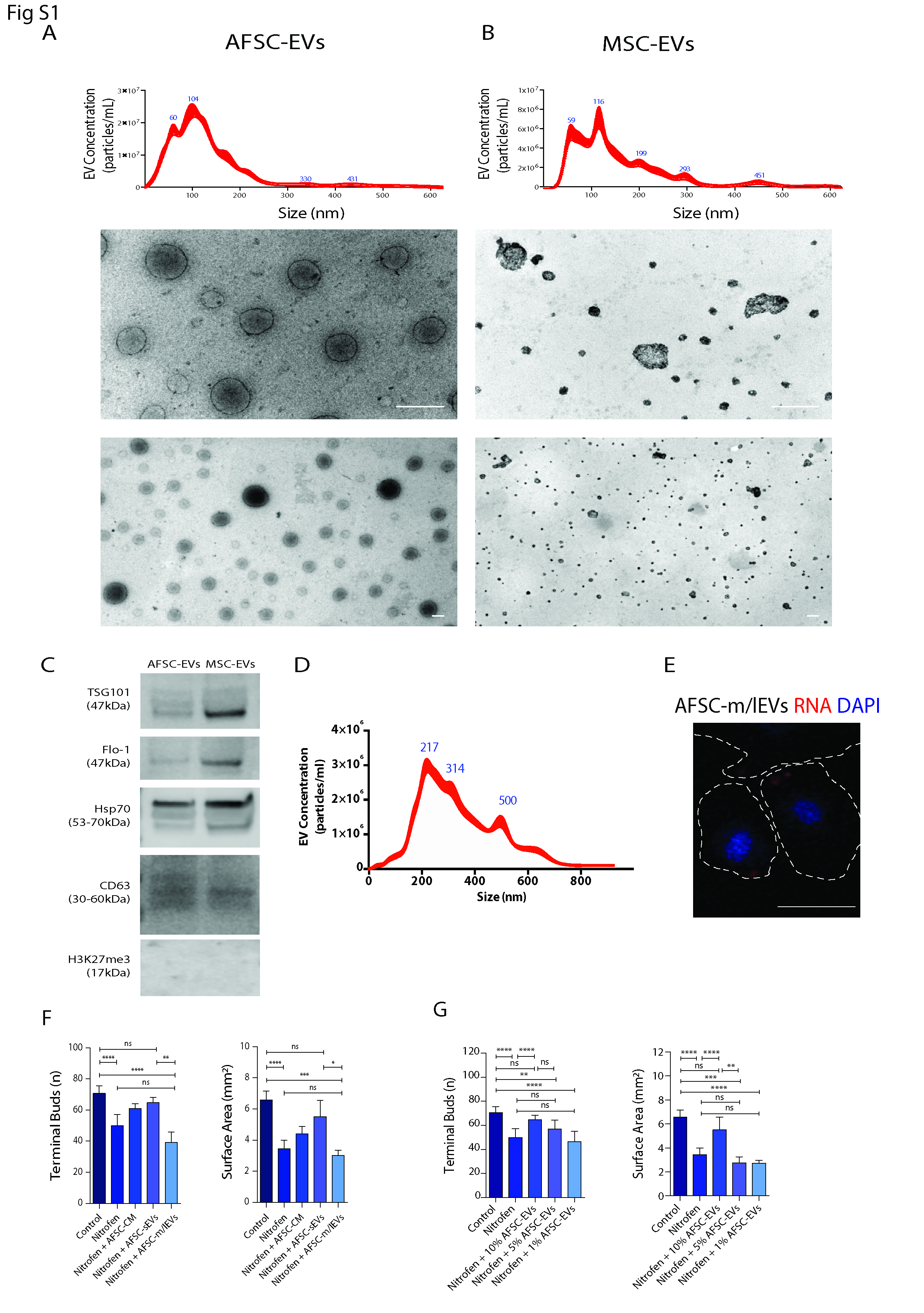

### bioRxiv-supp_Sup Fig 2.tif

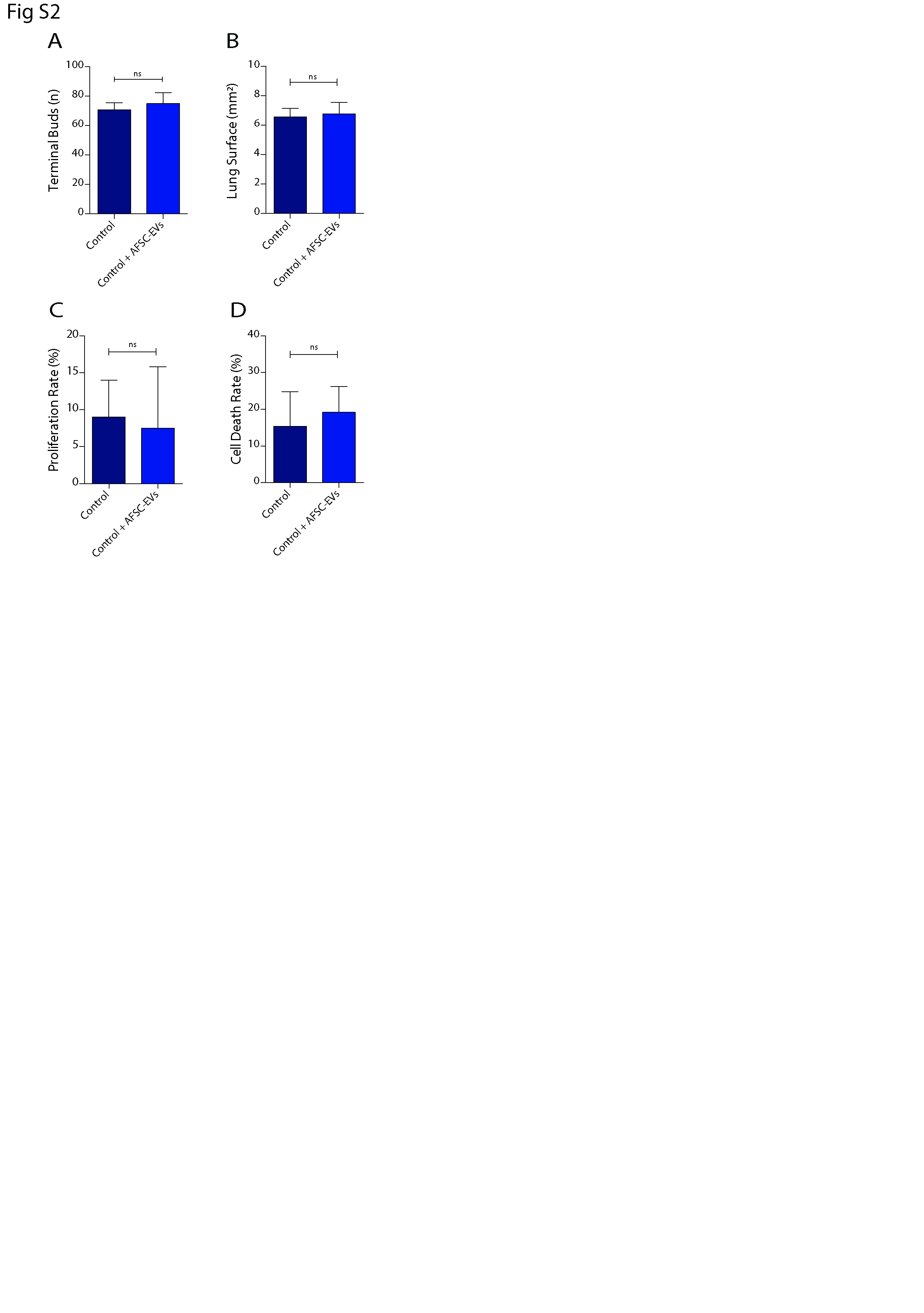

### bioRxiv-supp_Sup Fig 3.tif

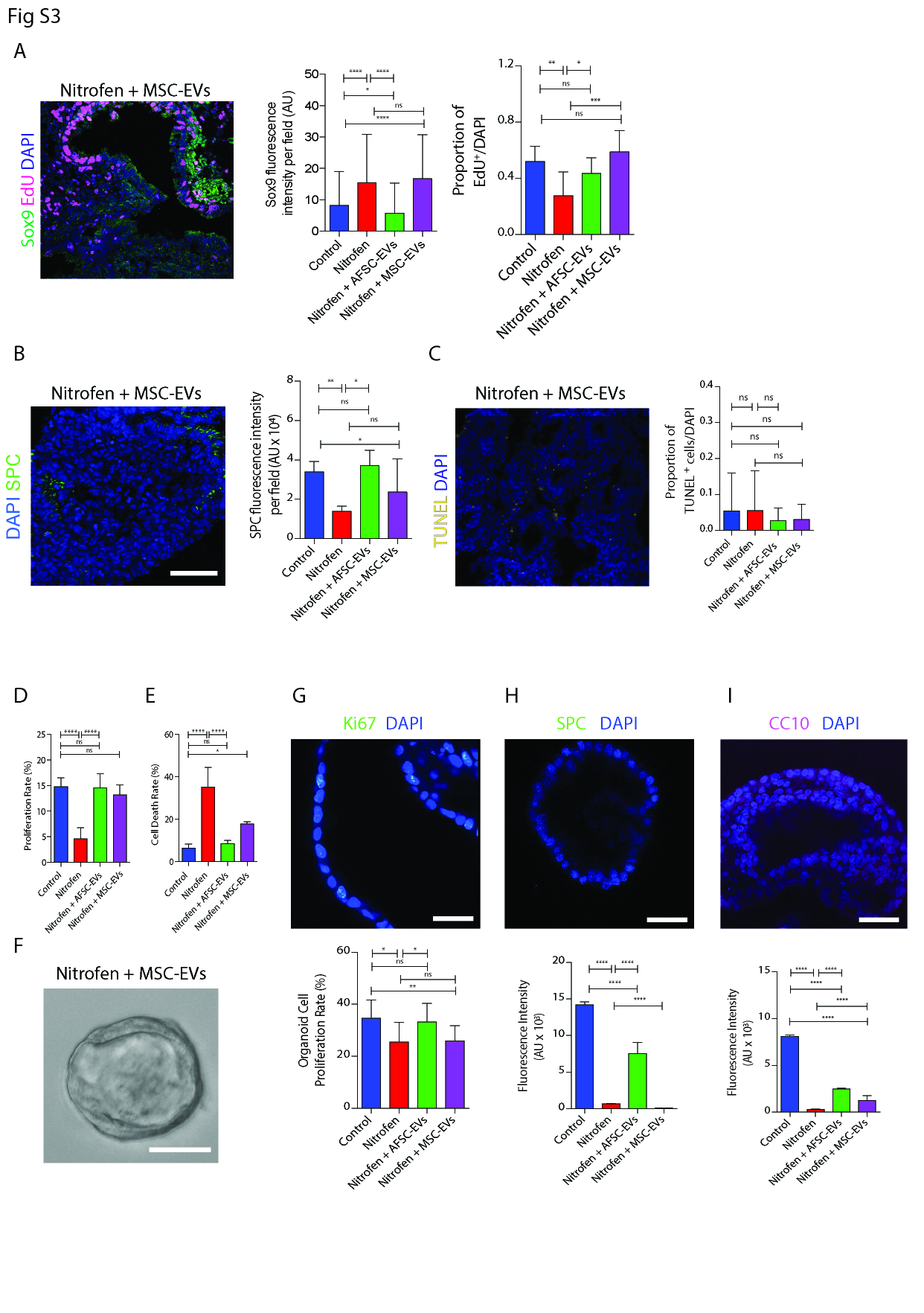

### bioRxiv-supp_Sup Fig 4.tif

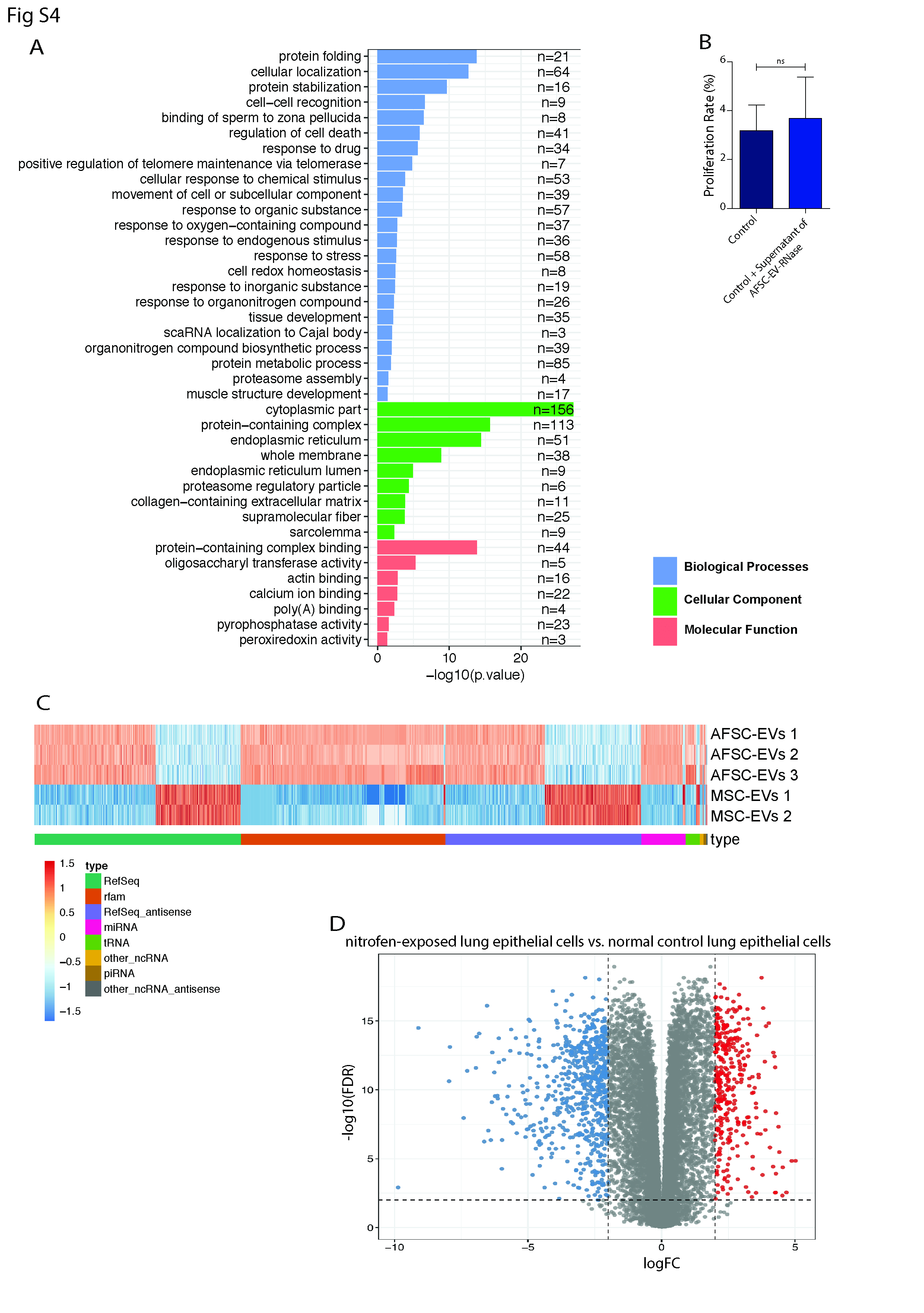

### bioRxiv-supp_Sup Fig 5.tif

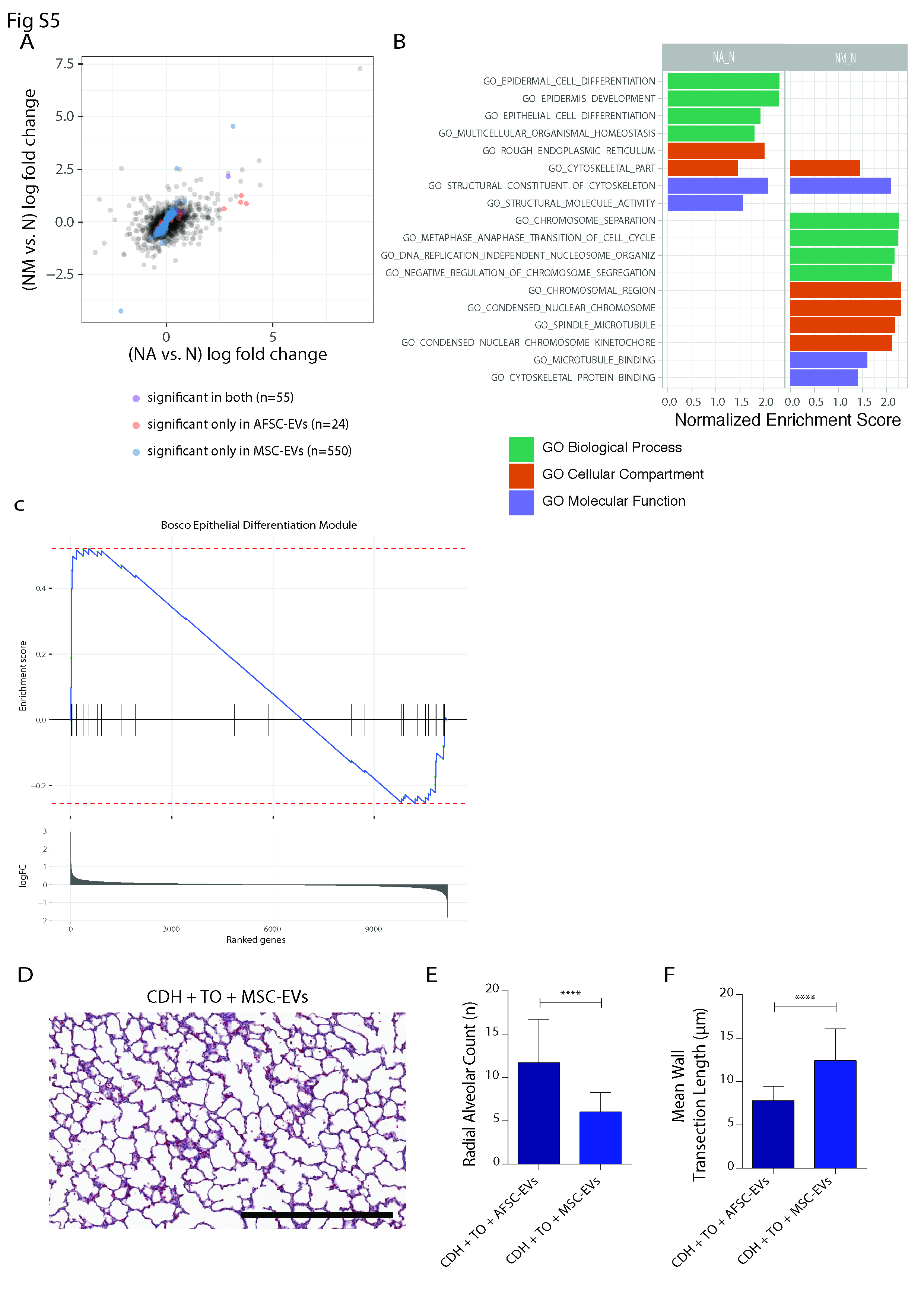

### bioRxiv-supp_Sup Fig 6.tif

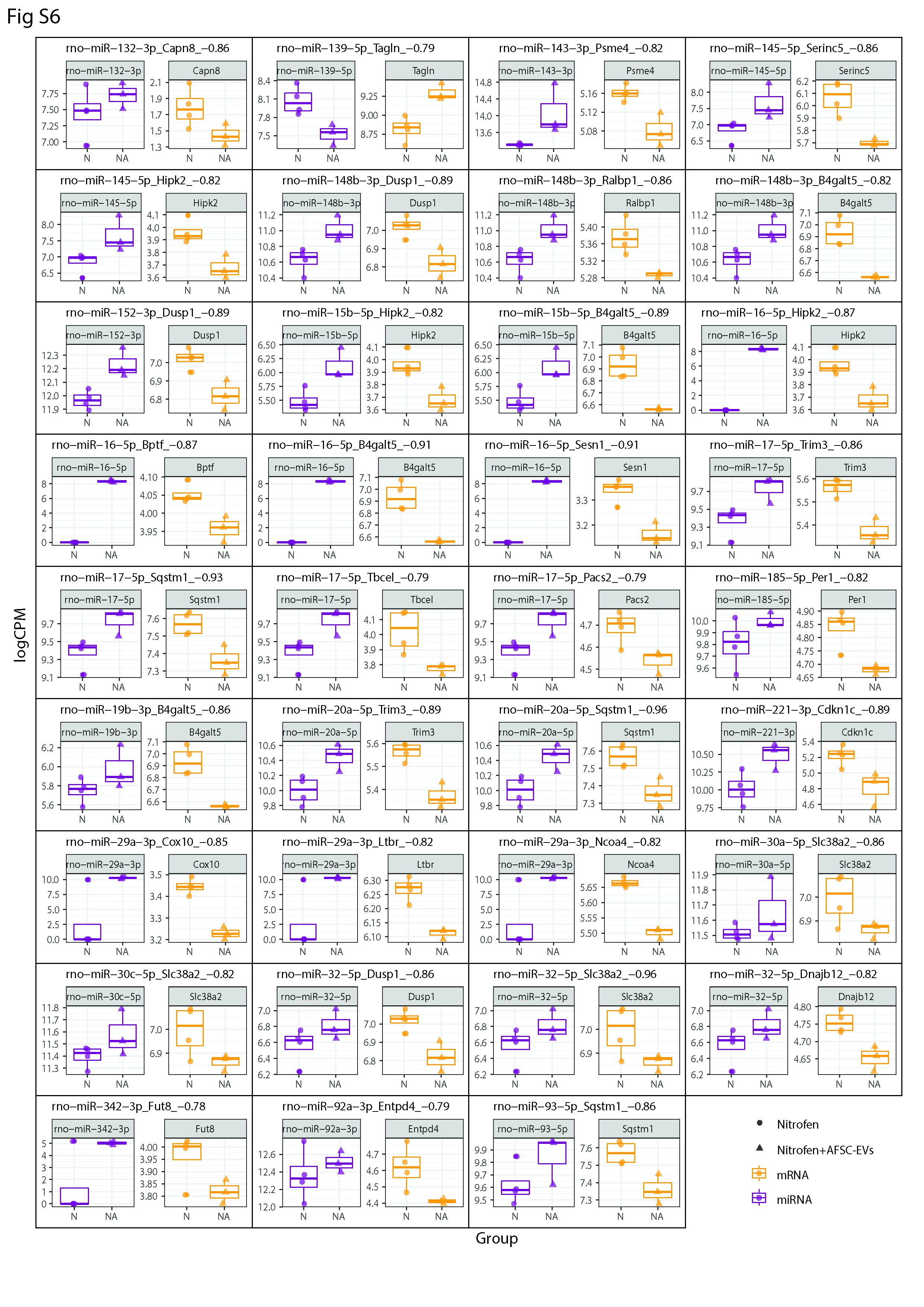

### bioRxiv-supp_Sup Fig 7.tif

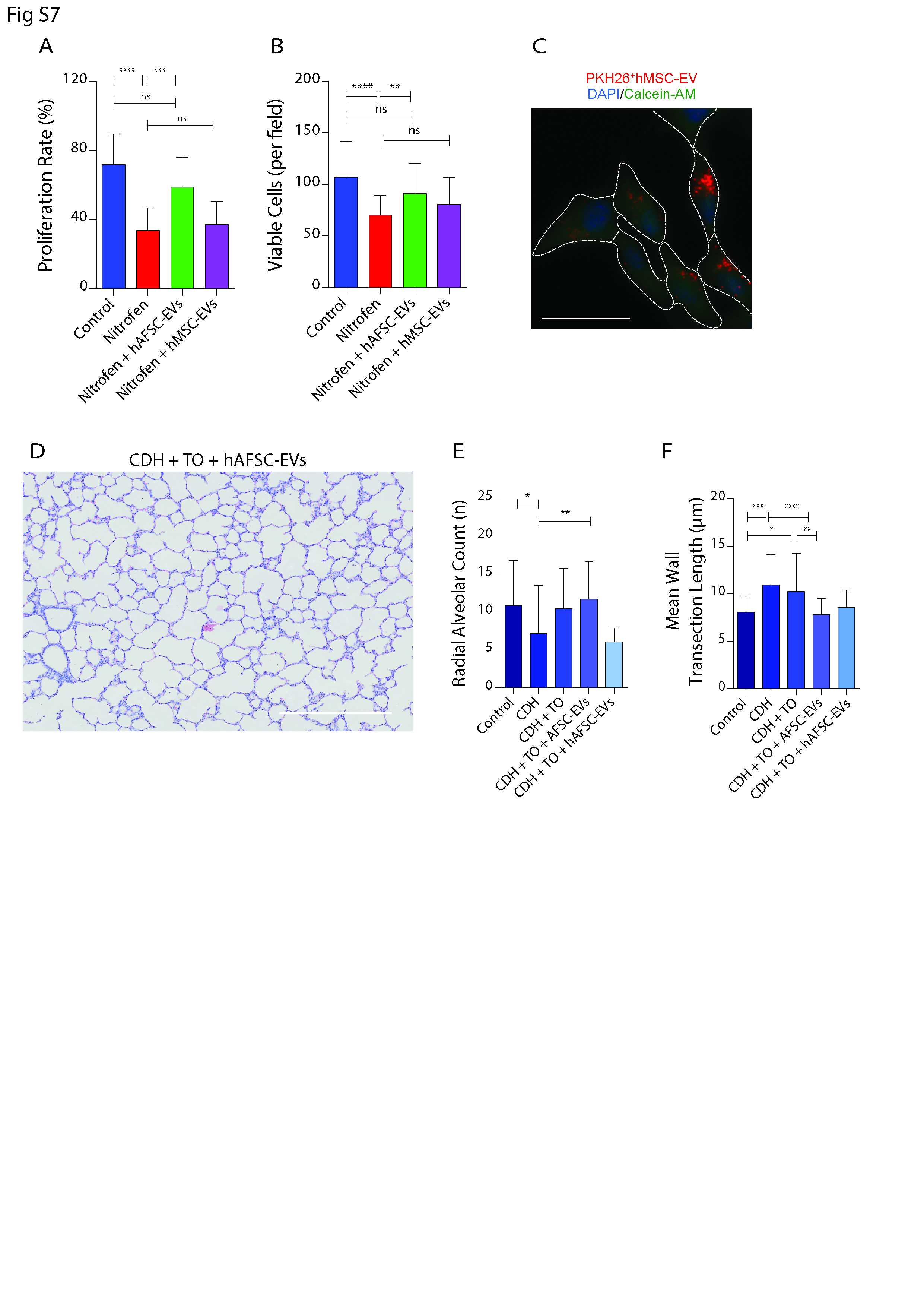

### bioRxiv-supp_Sup Fig 8.tif

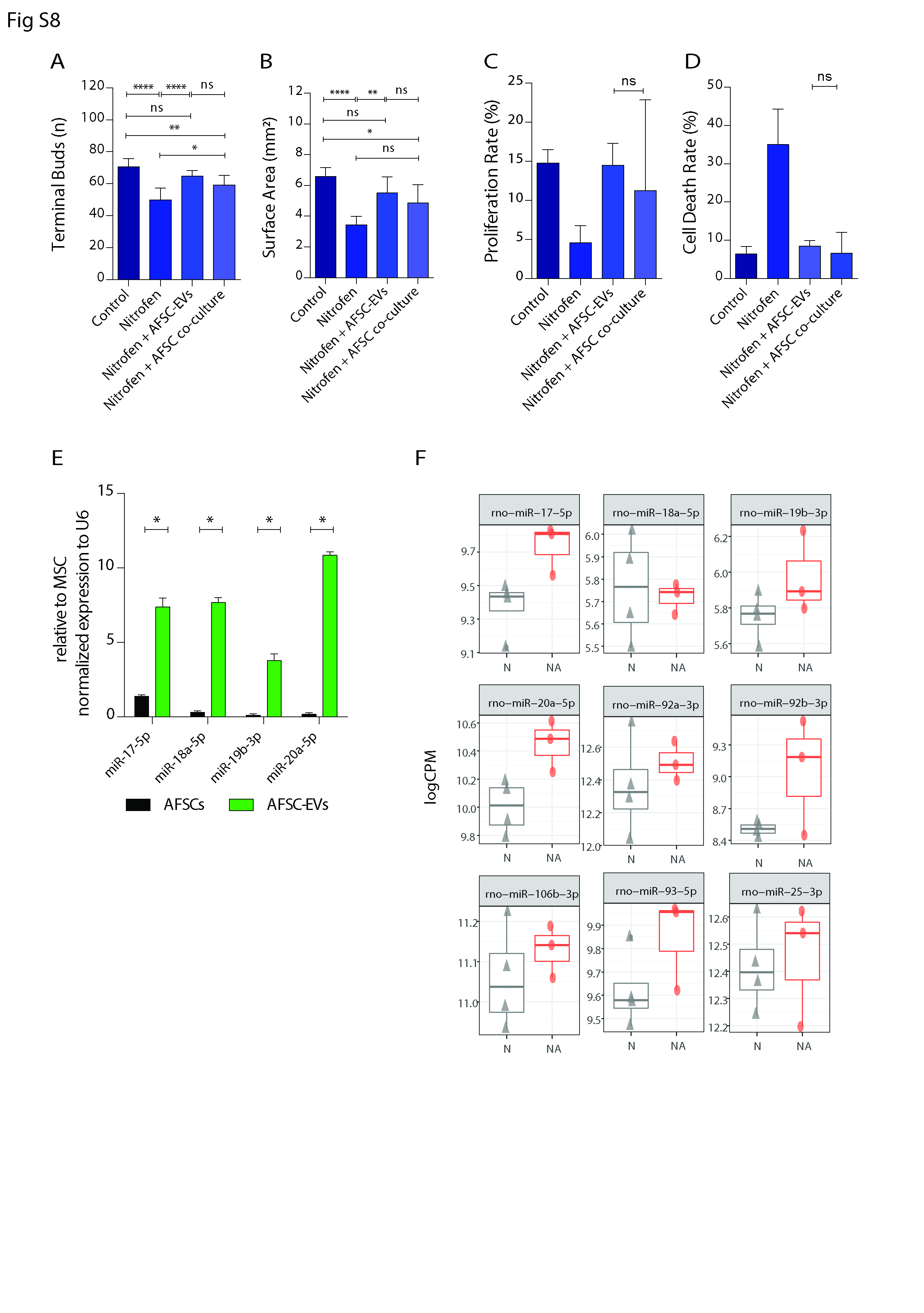
